# Discovery and biosynthesis of clostyrylpyrones from the obligate anaerobe *Clostridium roseum*

**DOI:** 10.1101/2020.08.10.245514

**Authors:** Jeffrey S Li, Yongle Du, Di Gu, Wenlong Cai, Allison Green, Samuel Ng, Alexander Leung, Antonio Del Rio Flores, Wenjun Zhang

## Abstract

Anaerobic bacteria are a promising new source for natural product discovery. Examination of extracts from the obligate anaerobe *Clostridium roseum* led to discovery of a new family of natural products, the clostyrylpyrones. The polyketide synthase-based biosynthetic mechanism of clostyrylpyrones is further proposed based on bioinformatic, gene knockout, biochemical analysis and heterologous expression studies.

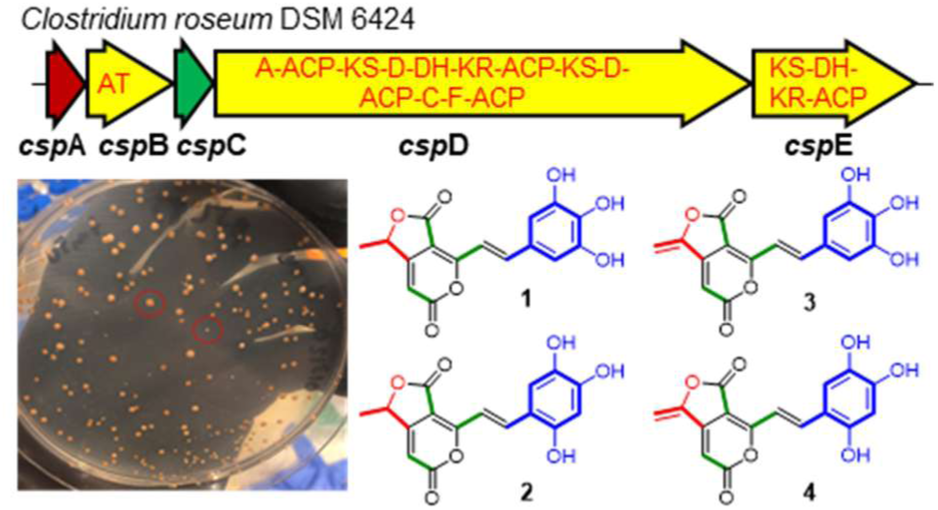

---

Natural products (NPs) are important sources of privileged chemical scaffolds for development of drugs and other bioactive compounds.^1^ Whereas natural product discovery has been historically biased toward more easily culturable aerobic microbes, anaerobes are now recognized as an underutilized reservoir for the discovery of NPs, based on genomic analyses demonstrating the widespread occurrence of biosynthetic gene clusters (BGCs) encoding major classes of NPs such as polyketides and non-ribosomal peptides.^2^ Increasingly available chemical data for anaerobe-derived NPs speak to the structural and functional diversity which can be accessed by exploring this resource.^3^–^8^

The Class Clostridia is a subset of the anaerobes which are particularly enriched in biosynthetic potential for polyketides based on genomic analyses.^9^ Polyketides are biosynthesized by characteristic multidomain assembly line enzymes which join malonyl-CoA monomers by decarboxylative Claisen condensation, often in iterative or modular processing.^10^ While polyketide synthase (PKS) genes have been detected in many Clostridia, very few examples of polyketides have been reported (**Figure 1**), all of which are associated with iterative PKSs. For example, clostrienose is a product of an iterative type I single-module PKS from *Clostridium acetobutylicum*.^11^ Clostrubins are aromatic polyketides biosynthesized by a type II PKS in both *Clostridium beijerinckii* and *Clostridium puniceum*.^12,13^ Through heterologous expression of a type II PKS BGC of *Blautia wexlerae* (belong to the class Clostridia) in a *Bacillus subtilis* host, a new aromatic polyketide, wexrubicin, was also discovered.^8^ These compounds exhibit structural and functional diversity and complexity, underscoring the value of studying polyketides from Clostridia.

**Figure 1.**
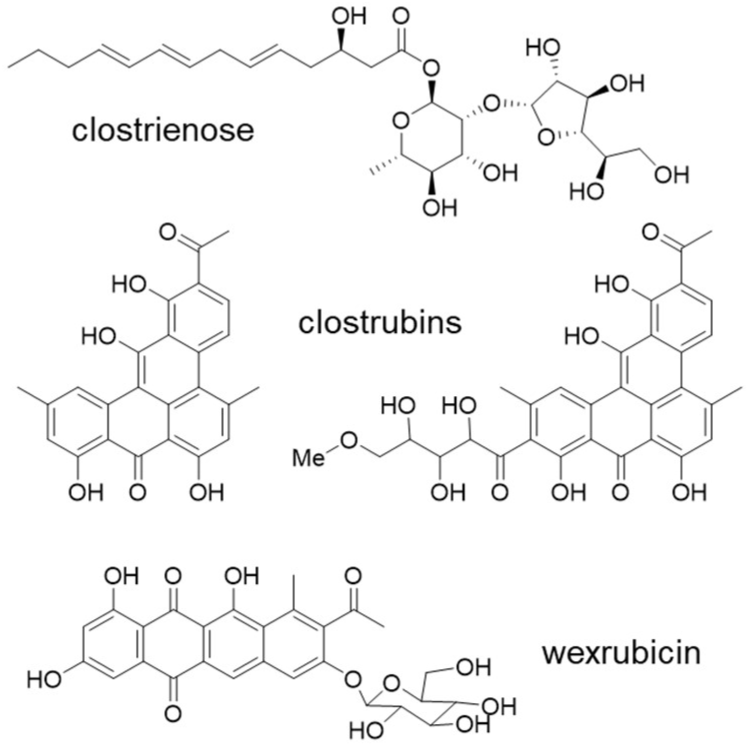
Polyketide NPs associated with Clostridia.

*Clostridium roseum* DSM 6424 (*Cro*), first isolated from German maize,^14^ is an important anaerobe with broad biotechnological significance for production of organic solvents and biofuels.^15^ Recently, *Cro* has been highlighted as one promising organism for NP discovery due to the presence of large thiotemplated BGCs in its genome.^3^ *Cro* is noteworthy among anaerobes in its production of pigments, including yellow-orange compounds which are secreted into the culture media. We thus firstly targeted these compounds for structural elucidation. High performance liquid chromatography (HPLC) examination of ethyl acetate extracts of *Cro* culture revealed the presence of several peaks with UV/vis maxima around 450 nm (**Figures 2** and **S1**). High resolution-electrospray ionization-mass spectrometry (HR-ESI-MS) analyses of the three major peaks (I-III) in positive ion mode enabled prediction of their associated chemical formulae: I, C_16_H_10_O_7_ (calculated for C_16_H_11_O_7_^+^ : m/z 315.0499; found: m/z 315.0503); II, C_16_H_12_O_7_ (calculated for C_16_H_13_O_7_^+^ : m/z 317.0656; found: m/z 317.0663); III, C_16_H_12_O_8_ (calculated for C_16_H_13_O_8_^+^ : m/z 333.0605; found: m/z 333.0607) (**Figure S1**). The predicted molecular formulae and similarities in the UV spectra suggested these peaks were congeners. For purification and structural elucidation, a four-liter batch culture of *Cro* was extracted and purified by size-exclusion chromatography with a Sephadex LH-20 column packing, followed by reverse-phase C18 HPLC purification. Peak I was isolated as a brownish amorphous solid with a yield of 0.4 mg; Peak II as an orange amorphous solid with 0.56 mg. Peak I material exhibited remarkable thermochromic and solvatochromic properties after isolation (**Figure S2**). The yield of Peak III was too low to be further characterized by NMR spectroscopic analysis.

**Figure 2.**
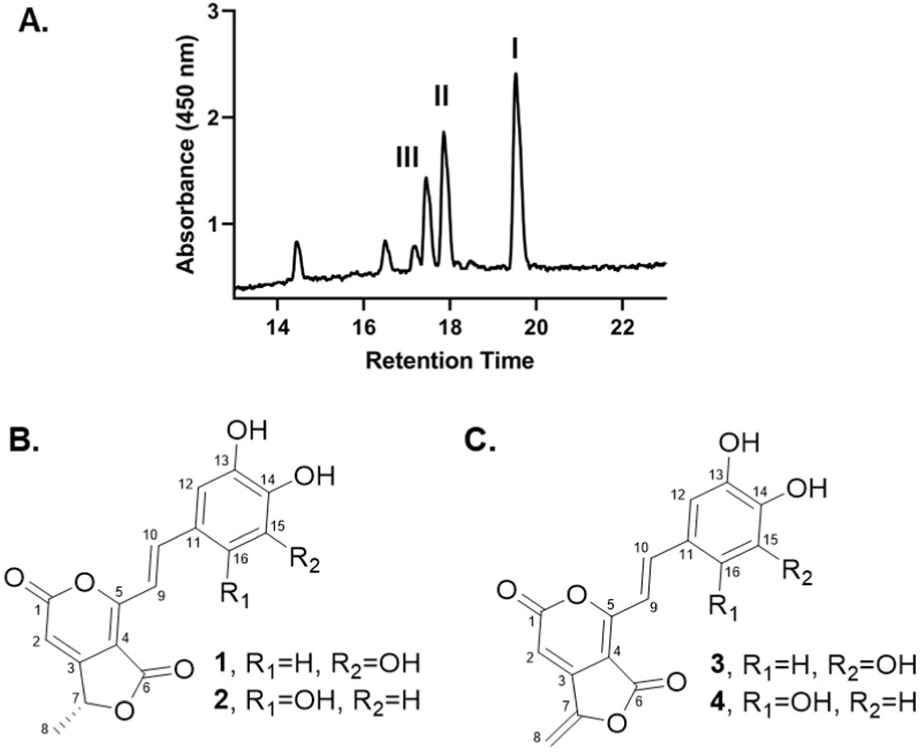
Discovery of clostyrylpyrones. (A) UV traces (450 nm) during LC-MS analysis of *Cro* extracts. (B) Structures of compounds isolated from Peak II. (C) Structures of compounds isolated from Peak I.

A detailed analysis of the 1D and 2D NMR spectra (^1^H, ^13^C, gHSQC, and gHMBC) of Peak I and II led to the discovery of four chemical structures, named the clostyrylpyrones A-D (**1-4**) (**Figures 2, S3** and **S4, Table S1**). In particular, Peak II contained a mixture of two structural isomers, **1** (59%) and **2** (41%), which were inseparable during HPLC purification; similarly, Peak I contained compounds **3** (53%) and **4** (47%). These deductions are supported by the following evidence: The obtained ^1^H NMR and HSQC spectra of peak II displayed signals for four hydroxyl groups (*δ*_H_ 9.34, 13/15-OH of **1**; *δ*_H_ 9.11, 13-OH of **2**; *δ*_H_ 9.08, 14-OH of **2**), two trans-disubstituted olefins (*δ*_H_ 7.62, H-10 of **2**; *δ*_H_ 7.55, H-10 of **1**; *δ*_H_ 7.30, H-9 of **1**; *δ*_H_ 7.27, H-9 of **2**), six aromatic or olefinic protons (*δ*_H_ 6.73, H-12 and H-16 of **1**; *δ*_H_ 6.70, H-12 of **2**; *δ*_H_ 6.58, H-15 of **2**; *δ*_H_ 6.30, H-2 of **1**; *δ*_H_ 6.23, H-2 of **2**), two oxygen bearing methines (*δ*_H_ 5.53, H-7 of **1/2**), and two methyl groups (*δ*_H_ 1.53, H_3_-8 of **1**/**2**). Furthermore, the ^13^C NMR and HSQC spectra showed signals for eighteen quaternary *sp*^2^ carbons, ten aromatic or olefinic methines, two oxygen bearing alkyl methines, and two methyl groups. Since all proton and carbon signals showed in pairs with similar chemical shifts, we proposed that peak II included two compounds with isomeric structures. The presence of each dilactone fragment was proposed through their proton and carbon chemical shifts and supported by their ^3^*J*-HMBC cross-peaks from H-2 to C-4, H-7 to C-2 and H-8 to C-3, as well as ^2^*J*-HMBC from H-2 to C-1 and H-7 to C-3. They were independently connected to a double bond based on ^3^*J*-HMBC correlations from H-9 to C-4 and H-10 to C-5. In addition, the other side of each double bond was linked to pyrogallol or hydroxyquinol by the observations from HMBC correlations. The absolute configuration of compounds **1/2** was assigned by the ECD spectrum (**Figure S5**). The n → π* transition bands at 243 nm showed a negative Cotton effect and 264 nm showed a positive Cotton effect, defining the absolute configuration at C-7 as R.^16,17^

The difference of 2 Da between the masses of **1**/**2** and **3**/**4** suggested that **3**/**4** might be dehydrogenated products of **1**/**2**. Dehydrogenation of the double bond of C-7 to C-8 in **1**/**2** was confirmed by the deshielding of the chemical shifts of C-7 and C-8 from *δ*_C_ 76.1 (CH) and 19.3 (CH_3_) in **1**/**2** to *δ*_C_ 150.7 (C) and 94.8 (CH_2_) in **3/4**, respectively. The NMR spectroscopic data of **3** and **4** were similar to those of **1** and **2** except as noted above and were fully consistent with its structural assignment as dehydrogenated products of **1** and **2** at C-7 and C-8, as confirmed by the HMBC cross-peaks (**Figure S4**).

We have thus discovered a group of new compounds sharing an unprecedented fused five- and six-membered dilactone scaffold with a trihydroxystyrene substituent at the C-5 position. It is notable that the styrene-substituted six-membered pyrone is a motif commonly found from eukaryotic phenylpropanoid NPs (**Figure S6**). These known styrylpyrones are often biosynthesized from aromatic amino acids which have undergone ammonia elimination and extension by type III PKS in plants^18,19^ or iterative type I PKS in fungi.^20^ We then reasoned that **1-4** could be biosynthesized by a similar logic, via a PKS promoted extension of an aromatic starter unit, although the enzymatic machinery for the assembly of the unique dilactone scaffold remained elusive.

To probe the biosynthesis of clostyrylpyrones, we scanned the *Cro* genome using antiSMASH^21^ for PKS-containing BGCs and identified a candidate locus for clostyrylpyrone biosynthesis, here designated as *cspA-E* (**Figure 3A**). Bioinformatic analyses of the five genes revealed a *trans*-AT five-module PKS complex, encoded by *cspB, cspD* and *cspE*, which is in agreement with the predicted five PKS building monomers of clostyrylpyrones (one aromatic starter unit and four extender units). *CspA* and *cspC* encode two modification enzymes, 3-de-oxy-D-arabinoheptulosonate 7-phosphate (DAHP) synthase and a dehydrogenase, which might be involved in the aromatic starter unit biosynthesis and PKS chain modification, respectively. To confirm the requirement of *csp* for clostyrylpyrone biosynthesis, a *cspD* inactivation mutant (Δ*cspD*) was constructed in *Cro* using a suicide vector through double crossover, and the genotype of the resulting mutant was verified by PCR (**Figure S7**). The production of clostyrylpyrones was completely abolished in Δ*cspD* (**Figure 4A**), confirming that the identified *csp* cluster is involved in the biosynthesis of clostyrylpyrones. No homologous gene clusters were identified from public databases based on multi-Blast analysis, consistent with the unique chemical structures of clostyrylpyrones.

**Figure 3.**
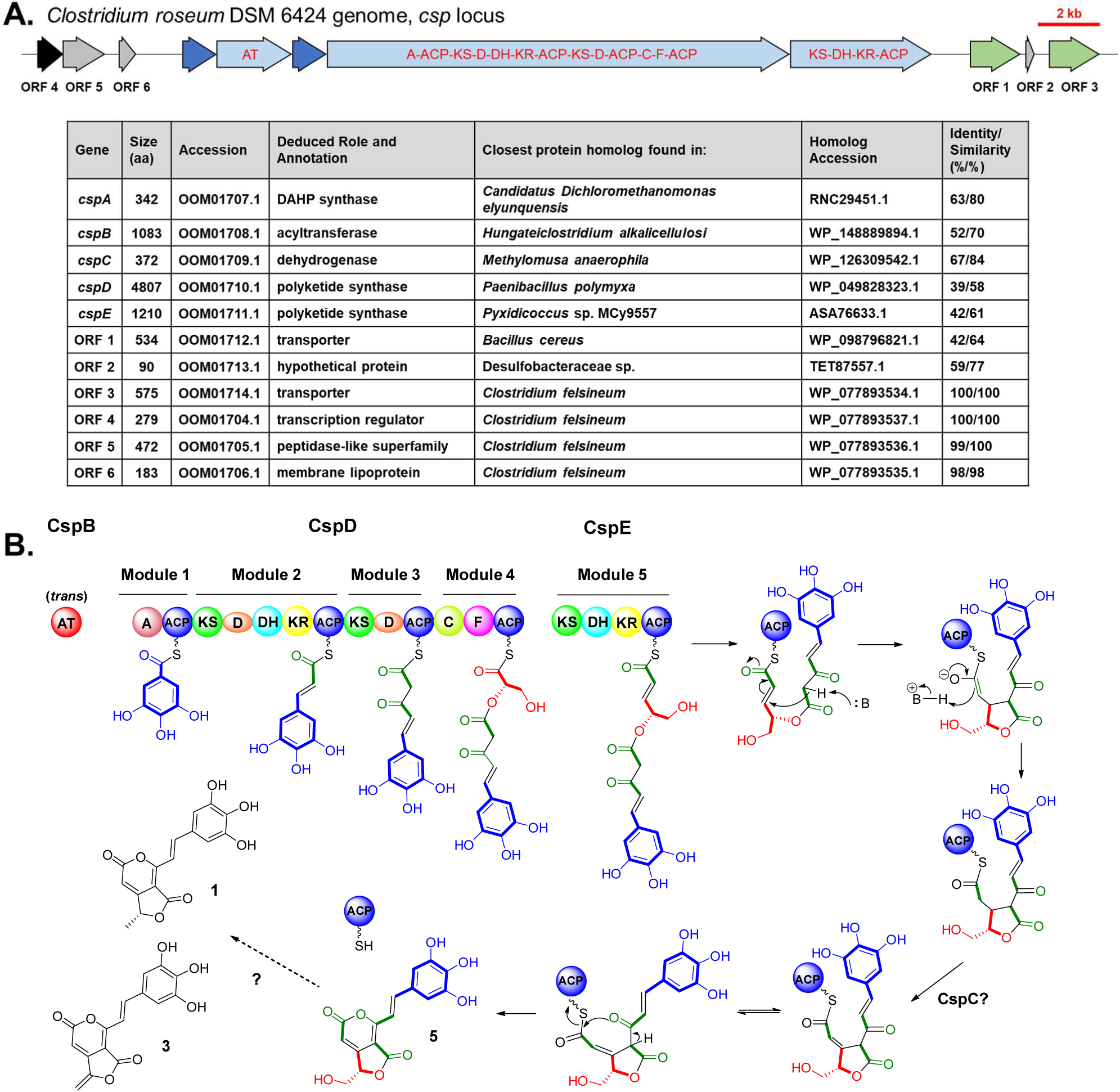
Map of *csp* gene neighborhood and proposed biosynthetic pathway. (A) Organization of *csp* genes, nearby open reading frames (ORFs), and their deduced roles based on sequence homology. Genes are colored by function: light blue, PKS related genes; dark blue, other biosynthetic genes; black, transcriptional regulator; grey, other; green, transporter. Domain abbreviations: AT, acyl-transferase; A, adenylation; ACP, acyl carrier protein; KS, ketosynthase; D, *trans*-AT docking; DH, dehydratase; KR, ketoreductase; F, FkbH-like. (B) Proposed biosynthesis of clostyrylpyrones. For simplicity, gallic acid is depicted as a representative starting monomer.

**Figure 4.**
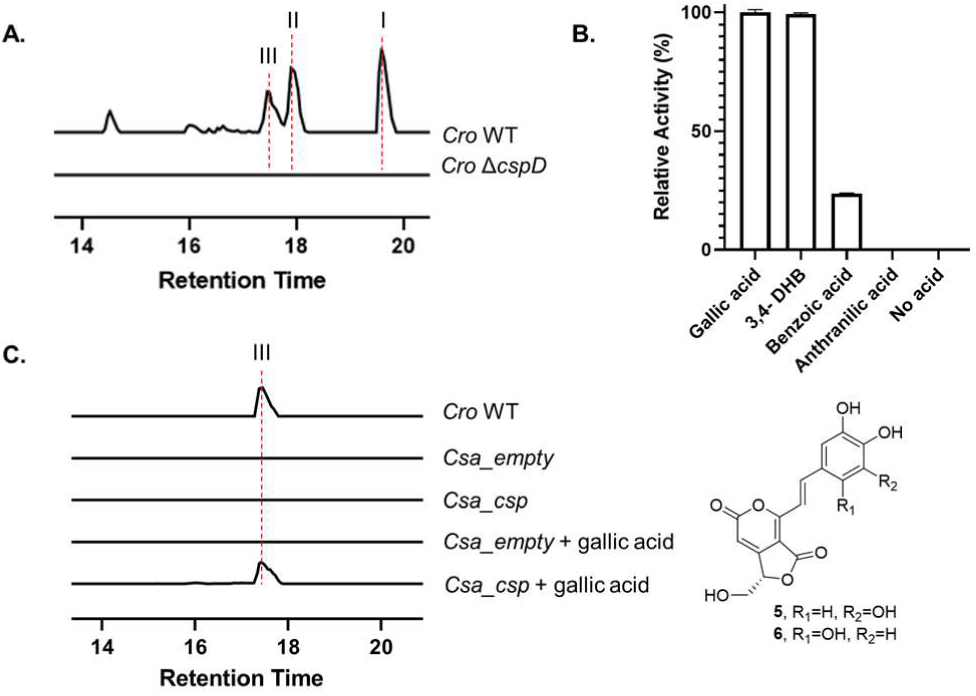
Biosynthetic study of *csp* products. (A) Extracted ion chromatograms showing deletion of *cspD* abolishes clostyrylpyrone production in *Cro*. The calculated masses (I: *m/z* = 315.0499 [M+H]^+^; II: *m/z* = 317.0656 [M+H]^+^; III: *m/z* = 333.0605 [M+H]^+^) with a 10-p.p.m. mass error tolerance were used. (B) Activity of the purified A-ACP didomain of CspD toward various acid substrates determined by ATP-PP_i_ release assays. Error bars represent standard deviations from at least three independently performed experiments. (C) Extracted ion chromatograms showing heterologous expression of *cspA-E*. Putative structures of compound **5**/**6** from peak III of *Cro* WT are also shown here.

The identification of *csp* has allowed us to propose a putative enzymatic pathway for clostyrylpyrone biosynthesis (**Figure 3B**). The PKS assembly line starts with the activation of 3,4,5-trihydroxybenzoic acid (gallic acid) or 2,4,5-trihydroxybenzoic acid by the initial adenylation (A) domain of CspD. *In silico* sequence alignment and modeling of this A domain identified residues potentially interacting with the aromatic substrates and yielded an unprecedented specificity motif^22^ (MPWGLVDGIK), consistent with incorporation of novel trihydroxybenzoic acid substrates (**Figure S8**). In addition, the activity of this A domain toward gallic acid activation has been biochemically confirmed using the purified A-ACP didomain of CspD (**Figure 4B and S9**). While the biogenesis of 2,4,5-trihydroxybenzoic acid remains unknown, gallic acid has been known to derive from the shikimic/chorismic acid pathways in both plant and bacterial model organisms.^23^ We further identified the predicted enzymes *in silico*, including the dedicated 3-dehydroshikimate reductase (AroE), involved in gallic acid biosynthesis which are encoded elsewhere on the *Cro* genome (**Figure S10)**. In addition, *cspA* encodes an extra copy of DAHP synthase, which catalyzes the committed first step in the shikimic/chorismic acid pathways. CspA is thus proposed to further channel the primary metabolites to the trihydroxybenzoic acid starter unit synthesis in *Cro*.

After the starter unit activation and loading onto CspD, the chain is elongated twice using malonyl-CoA by the next two typical PKS modules of CspD, before the action of the fourth atypical module of CspD with domains organized into C-F-ACP (C: condensation, F: FkbH-like domain, ACP: acyl carrier protein) (**Figure 3B)**. The presence of an F domain is reminiscent of the biosynthetic mechanism of agglomerin and other tetronate NPs which feature exocyclic-olefin bearing lactones, as in **3** and **4**.^24^ We thus propose that this module enables the incorporation of D-glycerate, derived from a primary metabolite such as D-1,3-bisphosphoglycerate, via a regiospecific ester bond formation into the PKS assembly line (**Figure S11**).^25,26^ After the last chain extension promoted by CspE, a Michael addition reaction takes place to generate the five-membered lactone, accompanied by the chain release from the assembly line through the six-membered pyrone formation. The PKS complex lacks an obvious chain release domain, and the fused five- and six-membered dilactone cyclization is thus most likely spontaneous associated with product release. CspC, a predicted dehydrogenase, may catalyze the double bond formation between C-2 and C-3 on or post the assembly line to yield the proposed hydroxylated intermediates, **5** and **6**. It is notable that **5/6** have chemical formulae of C_16_H_12_O_8_, consistent with the predicted formula associated with the Peak III minor product (**Figure 2** and **S12**). **3/4** could form from **5/6** through acetylation–elimination as observed in tetronate antibiotics biosynthesis,^27^ but the homologous acetyltransferase and eliminating enzyme were not identified from the genome of *Cro*, suggesting that the conversion of **5/6** to **3/4** and **1/2** may involve a different mechanism and the relevant biosynthetic genes are not clustered with *csp*.

To further confirm the role of *csp* in clostyrylpyrone biosynthesis, we cloned the 22 kb *cspA-E* into a plasmid for heterologous expression studies in *Clostridium saccharoperbutylacetonicum* N1-4 (*Csa*).^28^ Based on the proposed biosynthetic activity of *cspA-E*, we proposed that the functional expression of *csp* would yield product **5/6**. Since *Csa* lacks the ability to produce gallic acid (**Figure S10**), the culture of *Csa_csp* was supplemented with gallic acid for heterologous production. As expected, a new product was detected in the culture of engineered strain of *Csa_csp*. This product demonstrated identical retention time and mass-to-charge ratio to that of the Peak III minor product (predicted to be **5/6**) from the wild-type *Cro*, and the product was thus predicted to be **5** as its production required gallic acid (**Figure 4C**). The heterologous expression strain of *Csa_csp* did not produce peaks corresponding to compounds **1**-**4**, which is consistent with the prediction that *csp* lacks genes responsible for subsequent maturation of the biosynthetic intermediates **5/6**. This work thus adds another example of non-clustered biosynthetic genes in Clostridia, raising concerns toward employing a heterologous expression strategy for revealing native natural products.

Highly conjugated phenolic compounds are well known to be powerful antioxidants that can stabilize free-radical chain reactions in foods and other biochemical systems.^29^ For example, clostrubins have been reported to protect Clostridia from oxidative stress.^13^ Thus, we were interested in assessing the function of clostyrylpyrones as antioxidants in the context of *Cro*. A disc diffusion assay was performed to compare the ability of Cro WT and *Cro* Δ*cspD* in tolerating hydrogen peroxide stress. Despite different cellular morphology of the wide-type and mutant, the mutant appeared to be less tolerant toward hydrogen peroxide with an obvious inhibition zone, suggesting that clostyrylpyrones might be involved in coping with oxidative stress of anaerobes (**Figure S13**). Furthermore, we tested the cytotoxic activity of **1**/**2** against MCF-7, a human breast cancer cell line. The MTT-based tetrazolium staining assay^30^ was used to quantify cell viability and activity, and the growth rate inhibition (GR)^31^ was calculated and plotted (**Figure S13**). No obvious growth inhibitory activity was detected for **1**/**2**.

In summary, we here report the discovery of the clostyrylpyrones, a novel family of polyketides isolated from an obligate anaerobe. Structural elucidation is described for four compounds (**1** to **4**), all featuring an unprecedented scaffold of styrylpyrone fused to a tetronate-like heterocycle. We have further identified the biosynthetic gene cluster of clostyrylpyrones by genome mining, targeted gene deletion, biochemical analysis and heterologous expression, which sets the stage for further deciphering the chemical logic and enzymatic machinery in clostyrylpyrone biosynthesis. A complex modular PKS with a *trans*-acting AT are employed in biosynthesis, and the unique dilactone scaffold is formed due to the enzymatic activity of FkbH-like domain containing PKS module as well as the non-enzymatic cyclization promoted by Michael addition and pyrone formation. The discovery of the clostyrylpyrones expands the limited chemical space of known anaerobe-derived polyketides and encourages future efforts in mining new chemical scaffolds from anaerobes.

## ASSOCIATED CONTENT

### Supporting Information

The Supporting Information is available free of charge on the ACS Publications website.

Experimental details, NMR data, and Supplementary Figures 1-13.

## AUTHOR INFORMATION

### Notes

The authors declare no competing financial interest.

## ACKNOWLEDGMENTS

We thank J. Pelton (UC Berkeley) for helping with NMR spectroscopic analysis. This work was supported by the National Institutes of Health (DP2AT009148) and the Chan Zuckerberg Biohub Investigator Program.

