## Supporting Information for "Discovery and biosynthesis of clostyrylpyrones from the obligate anaerobe *Clostridium roseum*"

*Table of Contents*

**Pages S3-S7.** ***Supplementary Methods***

***Supplementary Tables***

**Page S8-S9.** ***Table S1.*** NMR spectroscopic data for compounds **1, 2** in DMSO-d6 and compounds **3, 4** in $CD_3OD$ (900 MHz)

**Page S10.** ***Table S2.*** Strains and plasmids used in this work

**Page S11.** ***Table S3.*** Synthetic oligonucleotides used in this work

***Supplementary Figures***

**Page S12.** ***Figure S1.*** UV and HRMS characterization of clostyrylpyrones

**Page S13.** ***Figure S2.*** Chromism of **3** and **4**

**Page S14-S17.** ***Figure S3.*** NMR characterization of **1**, **2**

**Page S18-S20.** ***Figure S4.*** NMR characterization of **3**, **4**

**Page S21.** ***Figure S5.*** ECD spectrum analysis of compounds **1** and **2**

**Page S22.** ***Figure S6.*** Selected styrylpyrone NPs from fungi and plants

**Page S23.** ***Figure S7.*** Generation of *Cro* Δ*cspD.*

**Page S24-S25.** ***Figure S8.*** *In silico* analysis of CspD.

**Page S26.** ***Figure S9.*** Biochemical analysis of A-ACP didomain of CspD

**Page S27.** ***Figure S10.*** Proposed pathway of gallic acid biosynthesis in *Cro*

**Page S28*.*** ***Figure S11.*** The activity of FkbH-like domain

**Page S29.** ***Figure S12.*** MS/MS spectra of peak I, II and III

**Page S30.** ***Figure S13*.** Bioactivity assays of clostyrylpyrones

***References***

***Supplementary Methods***

**Bacterial strains and growth conditions**. All strains used in this study are listed (**Table S2**). *E. coli* strains were cultured in lysogeny broth (LB) at 37°C and supplemented with antibiotics when appropriate. Cloning was performed in *Escherichia coli* XL1-blue. *Csa* was maintained in PL7 media[1] (30 g/liter glucose, 5 g/liter yeast extract, 2.67 g/liter ammonium sulfate, 1 g/liter NaCl, 0.75 g/liter monobasic sodium phosphate, 0.75 g/liter dibasic sodium phosphate, 0.5 g/liter cysteine-HCl monohydrate, 0.7 g/liter magnesium sulfate heptahydrate, 20 mg/liter manganese sulfate monohydrate, and 20 mg/liter iron sulfate heptahydrate, with the initial pH adjusted to 6.5 using 1 N HCl). *Cro* was maintained at 30°C in an anaerobic chamber (Coy Laboratory Products, Grass Lake, MI) containing an atmosphere of 97% nitrogen and 3% hydrogen. For routine culture it was incubated in CBM containing 30 g/liter glucose, 0.5 g/liter monobasic potassium phosphate, 0.5 g/liter dibasic potassium phosphate, 4 g/liter tryptone, 0.2 g/liter magnesium sulfate heptahydrate, 10 mg/liter manganese sulfate heptahydrate, 10 mg/liter ferrous sulfate heptahydrate, 1 mg/liter para-aminobenzoic acid, 1 mg/L thiamine hydrochloride, and 2 µg/liter biotin with the pH adjusted to 6.5 with 1 N HCl.[2] The media was prepared as a filter-sterilized 2x concentrate and mixed with either water or 3% agar for liquid or solid media. For long-term storage, cultures were kept at -80°C in 20% glycerol.

**Plasmid construction.** Synthetic oligonucleotides were provided by IDT (Coralville, IA). The pMTL modular shuttle vector series were obtained from Chain Biotech (United Kingdom).[3] All sequences are in the 5' to 3' orientation. Phusion polymerase was used for all PCR reactions. FastDigest restriction enzymes were obtained from ThermoFisher. FseI and AscI were obtained from NEB. Plasmid constructs (**Table S2**) were assembled by the method of Gibson[4] and reaction mixtures were transformed into chemically competent *E. coli* XL1-blue. Clones were isolated and DNA was extracted using a Zyppy Plasmid Miniprep Kit (Zymo Research, Irvine, CA). Constructs were validated by restriction digest patterning and Sanger sequencing.

The *csp* suicide vector pJL83 was constructed as follows. pMTL82151 was digested by a cocktail of AscI, PmeI, and FseI. The 0.8 and 2 kb bands corresponding to *catP* (the chloramphenicol/thiamphenicol resistance gene cassette) and the Gram-negative origin of replication, respectively, were isolated. The upstream and downstream homology regions were amplified from purified *Cro* genomic DNA by primers JL322 & JL323 and JL324 & JL325 respectively.

The *csp* heterologous expression vector pJL109, based on pMTL83353 was constructed as follows. The intermediate vector pSN4 was constructed from the 3.8 kb plasmid backbone after restriction digestion by FseI and PmeI, and a PCR product amplifying the pWIS-empty vector[1] erythromycin resistance cassette via primers SN7 & SN8. Then, the pSN4 was linearized at the multiple cloning site by digestion with NotI and HindIII. The 22 kb *csp* locus was amplified in three overlapping pieces from purified *Cro* genomic DNA using JL401 & JL406, JL409 & JL410, and JL407 & JL402. The P-bdh promoter was amplified from the *Csa* genome using JL399 & JL400, and joined by splicing by overlap extension PCR to the PCR product containing the 5' sequence of *csp*, using the JL399 & JL406 primers. This enabled assembly of the full vector as a 4-piece reaction.

**Conjugation procedure**. Chemically competent *E. coli* WM6026 was freshly transformed with plasmid to serve as the conjugation donor. From this, overnight cultures of donor were incubated in LB with diaminopimelic acid (DAP) and 25 µg/ml chloramphenicol at 37°C. *Cro* wild-type overnight culture was prepared in PL7 at 30°C. At ~20 h, *Cro* cultures reached $OD_{600}$ 0.6-1. *E. coli* donors were washed twice in LB and transferred into the anaerobic chamber as a cell pellet. Pellets were resuspended in the PL7 conjugation recipient cultures, to concentrate donor to $OD_{600}$ of 6. Donor/recipient mixtures were plated (100 µl) onto CBM agar and incubated at 30°C for 3-8 h (optimally 4 h) before overlaying with 2.5 mg thiamphenicol in aqueous solution containing 10% DMSO. Conjugant colonies appeared after 1-3 days.

**Isolation of knockout mutants**. Knockout vectors were introduced into *Cro* by conjugation from *E. coli* WM6026. After 3 days, colonies were picked and cultured in 10 ml CBM and 100 µg/ml thiamphenicol. Liquid media was supplemented with 0.1% tween 80 to aid in culture dispersion. After 24 h, the culture was vortexed to homogenize, passaged at 0.1% into 10 ml subculture, and spread onto CBM agar supplemented with thiamphenicol. After 2-3 days, colonies were screened by touchdown PCR using Phusion polymerase. After isolation, mutants were cured of plasmid by 1-day culture in non-selective CBM followed by plating on CBM agar. Colonies were screened by PCR to confirm plasmid loss.

**Metabolomic analysis.** Metabolomics samples were analyzed in biological quadruplicate using LC-UV-HRMS. Chemical extracts were prepared by 1:1 extraction of 1 ml culture with 1 ml ethyl acetate. Mixtures were vortexed and spun down (6000×g, 1 min). The upper phase solvent layer was pipetted into a centrifuge tube and dried under $N_2$, then resuspended in 100 µl methanol. Injections (5 µl) were analyzed on an Agilent Technologies 6545 Accurate-Mass QTOF LC-MS instrument fitted with a 1290 Infinity II DAD for UV/vis and an Agilent Eclipse Plus C18 column (4.6×100 mm). The run method used a linear gradient of 2-98% $CH_3CN$ (v/v) over 40 min in $H_2O$ with 0.1% formic acid (v/v) at a flow rate of 0.5 ml/min. Data analysis was performed in MS-DIAL.[5]

**Purification, NMR and ECD characterization of clostyrylpyrones.** *Cro* inoculum was prepared in CBM (20 ml). After 2 days at 30°C, the densely grown, lightly pigmented cultures were aliquoted into two 2-liter bottles of freshly prepared CBM. Cultures were incubated without agitation for 5 days before harvesting. Culture was extracted twice with 1 volume ethyl acetate and gently centrifuged (2000×g, 10 min) to facilitate phase separation. The upper solvent layer was collected and removed under reduced pressure. The brown oily residue was suspended in 15 ml methanol and partially re-dissolved, leaving behind a red precipitate. The extract was clarified by centrifugation and loaded onto a size-exclusion column packed with Sephadex LH-20 (Sigma-Aldrich) and manually fractionated using MeOH mobile phase. Fractions were screened by LC-UV-MS using an Agilent Technologies 6120 Quadrupole instrument with a 1260 series DAD and Agilent Eclipse Plus C18 column (4.6×100 mm). The run method used a linear gradient of 2-98% $CH_3CN$ (v/v) over 15 min in $H_2O$ with 0.1% formic acid (v/v) at a flow rate of 0.5 ml/min. Fractions containing compounds **1** and **2** or **3** and **4** were consolidated, concentrated under vacuum, and purified using reverse-phase HPLC (using an Agilent 1260 HPLC with DAD) fitted with a semi-preparative Phenomenex Luna C18 column (5 µm, 10×250 mm, 100Å). Compounds **1** and **2** products were resolved with an isocratic elution in 28% $CH_3CN$ (v/v). Compounds **3** and **4** products were resolved with an isocratic elution in 30% $CH_3CN$ (v/v), while they were very unstable compared to **1**/**2**, and they decomposed rapidly (half life of ~1 day in methanol or DMSO) during purification and NMR experiments. All NMR spectra of the isolated compounds were acquired with a 900 MHz Bruker Avance III NMR spectrometer ($^1H$: 900 MHz, $^{13}C$: 226 MHz) equipped with a cryoprobe. The ECD spectrum of compound **1**/**2** was acquired with a Jasco J-815 Circular Dichroism (CD) Spectropolarimeter.

**Heterologous expression of *csp*.** The *csp* heterologous expression construct (pJL109) and an empty vector (pWIS_empty) were introduced into *Csa* by a previously described electroporation method.[6] Transformants of the two strains were cultured in 10 ml PL7 + 40 µg/ml erythromycin. Cultures were incubated at 30°C, with or without the addition of 1 mM gallic acid (Fisher), added from a 100× stock solution in ethanol. After 3 days, 1 ml samples were collected for metabolomic analysis.

**Construction of pDG16 for protein expression in *E. coli*.** The A-ACP didomain gene was amplified using pJL109 as a template (**Table S2**). The vector was constructed using the aLICator LIC Cloning and Expression Kit 4 containing an N-terminal His-tag and WQ site (Thermo) according to the manufacturer's protocol and instructions. The primers used to construct pDG16 are listed in **Table S3.** Plasmids were isolated using a Zyppy Miniprep Kit (Zymo Research) and confirmed by DNA sequencing (UC Berkeley DNA Sequencing Facility).

**Overexpression of A-ACP didomain of CspD.**

The expression and purification of A-ACP didomain of CspD followed the same general procedure for $His_6$-tag purification as detailed here. BAP-1 chemical competent cells were grown at 37°C in 1L of LB broth in a shake flask supplemented with 100 µg/mL of ampicillin to an OD600 of 0.5 at 200 rpm. The shake flask was then placed on ice for 10 mins and induced with 120 µM of isopropyl-β-D-thiogalacto-pyranoside (IPTG). The cells were incubated for 16 hours at 16°C and 200 rpm to undergo protein expression. Subsequently, the cells were harvested by centrifugation (6,000 x g, 15 min, 4°C), and the supernatant was removed. The cell pellet was resuspended in 30 mL of lysis buffer (25 mM HEPES pH 8, 500 mM NaCl, 5 mM imidazole) and cells were lysed by sonication on ice. Cellular debris was removed by centrifugation (15,000 x g, 30 min, 4°C) and the supernatant was filtered with a 0.45 µm filter before batch binding. Ni-NTA resin (Qiagen) was added to the filtrate at 2 mL/L of cell culture, and the samples were nutated for 1 hour at 4°C. The protein-resin mixture was loaded onto a gravity flow column. The flow through was discarded and the column was then washed with approximately 35 mL of wash buffer (25 mM HEPES pH 8, 100 mM NaCl, 20 mM imidazole) and tagged protein was eluted in approximately 22 mL of elution buffer (25 mM HEPES pH 8, 100 mM NaCl, 250 mM imidazole). The complete process was monitored using a Bradford assay. The purified protein was concentrated and exchanged into wash buffer (25 mM HEPES pH 8, 100 mL NaCl) using 30,000 MWCO Amicon ultra filter units. After two rounds of exchange, glycerol was added to the purified protein to a final concentration of 10%. The protein was flash frozen in liquid nitrogen and stored as beads at -80 ◦C. The presence and purity of the protein was assessed using SDS-PAGE and the concentration was determined using a NanoDrop UV-Vis spectrophotometer (Thermo Fisher). The approximate protein yield was 8.75 mg/L (67.1 kDa). This protein was used to demonstrate adenylation of gallic acid and other substrates in vitro.

**ATP-$PP_i$ release assays to determine relative activity of A-ACP didomain of CspD to Gallic Acid and other Substrates.**

The inorganic pyrophosphate released by the A-ACP didomain of CspD enzymatic reaction was measured continuously using the EnzChek Pyrophosphate Assay Kit (Thermo Fisher). A typical 100 µL assay contained 50 mM HEPES pH 8, 5 mM acid substrate, 5 mM ATP, 2 mM $MgCl_2$, and 5 µM A-ACP didomain of CspD. The MESG substrate, purine nucleoside phosphorylase and inorganic pyrophosphatase were added according to the protocol. Reactions were initiated by the addition of the acid substrate and monitored at 360 nm using a SpectraMax M2 plate reader. Initial velocities and the relative activity of each acid substrate with respect to gallic acid was calculated.

**Peroxide disc diffusion assay.** Assays were based on a reported method.[7] *Cro* wild-type and Δcsp CBM liquid cultures were inoculated from plates. At 18 h the cultures were turbid and $OD_{600}$ was diluted to 0.2 and 100 µl was plated on CBM to form a lawn. Filter discs of 6 mm diameter (GE Healthcare, Chicago, IL) were placed on top of the agar and dilutions of hydrogen peroxide solution were spotted at 10 µl. After 3 days, plates were examined for inhibition zones.

**Growth rate inhibitory assay**. Human MCF-7 cell cultures were obtained from the Berkeley Cell Culture Facility. Cell culture and quantification was performed using the MTT Cell Growth Assay Kit (Sigma Aldrich) using the recommended procedure.

***Table S1.*** NMR spectroscopic data for compounds **1**, **2** in DMSO-$d_6$ (900 MHz)

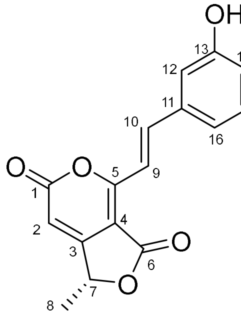


**1**, clostyrylpyrone A

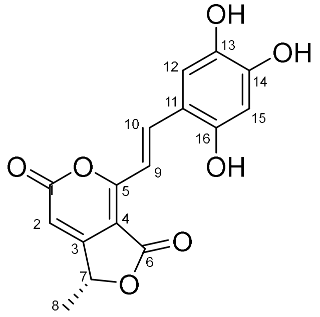


**2**, clostyrylpyrone B

| | **1** | | | **2** | | |
|---|---|---|---|---|---|---|
| position | $\delta_H$ ($J$ in Hz) | $\delta_C$, type | HMBC | $\delta_H$ ($J$ in Hz) | $\delta_C$, type | HMBC |
| 1 | | 159.7, C | | | 159.9, C | |
| 2 | 6.30 s | 101.8, CH | 1, 4, 7 | 6.23 s | 100.7, CH | 1, 4, 7 |
| 3 | | 163.6, C | | | 163.9, C | |
| 4 | | 102.9, C | | | 102.0, C | |
| 5 | | 162.7, C | | | 163.2, C | |
| 6 | | 166.4, C | | | 166.5, C | |
| 7 | 5.53 q (6.8) | 76.1, CH | 2, 3, 8 | 5.53 q (6.8) | 75.9, CH | 2, 8 |
| 8 | 1.53 d (6.8) | 19.3, $CH_3$ | 3, 7 | 1.53 d (6.8) | 19.3, $CH_3$ | 3, 7 |
| 9 | 7.30 d (15.6) | 110.3, CH | 4, 5, 11 | 7.27 d (15.6) | 108.7, CH | 4, 5, 11 |
| 10 | 7.55 d (15.6) | 142.3, CH | 5, 12/16 | 7.62 d (15.6) | 143.9, CH | 5, 12, 15 |
| 11 | | 124.8, C | | | 124.4, C | |
| 12 | 6.73 s | 108.1, CH | 10, 14, 16 | 6.70 s | 110.7, CH | 10, 14, 15 |
| 13 | | 146.4, C | | | 141.0, C | |
| 14 | | 138.0, C | | | 143.5, C | |
| 15 | | 146.4, C | | 6.58 s | 101.4, CH | 12, 14, 16 |
| 16 | 6.73 s | 108.1, CH | 10, 12, 14, 15 | | 152.8, C | |
| 13-OH | 9.34 br s | | 12, 13, 14 | 9.11 s | | 12, 14 |
| 14-OH | 9.08 s | | 13/15 | | | |
| 15-OH | 9.34 br s | | 14, 15, 16 | | | |

***Table S1 (continued).*** NMR spectroscopic data for compounds **3**, **4** in $CD_3OD$ (900 MHz)

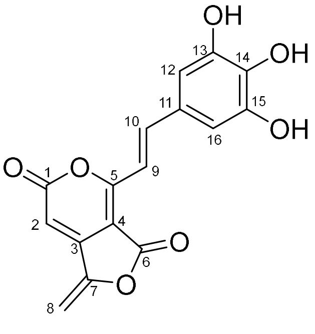


**3**, clostyrylpyrone C

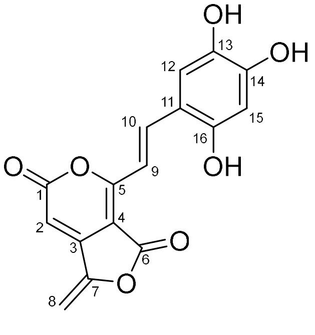


**4**, clostyrylpyrone D

| | **3** | | | **4** | | |
|---|---|---|---|---|---|---|
| position | $\delta_H$ ($J$ in Hz) | $\delta_C$, type | HMBC | $\delta_H$ ($J$ in Hz) | $\delta_C$, type | HMBC |
| 1 | | n.d.[a] | | | 160.8, C | |
| 2 | 6.39 s | 97.9, CH | 4 | 6.50 s | 99.6, CH | 1, 4 |
| 3 | | n.d.[a] | | | n.d.[a] | |
| 4 | | 99.7, C | | | 101.4, C | |
| 5 | | 164.8, C | | | 164.0, C | |
| 6 | | n.d.[a] | | | n.d.[a] | |
| 7 | | 150.7, C | | | n.d.[a] | |
| 8 | 5.52 d (3.5) | 94.8, $CH_2$ | 7 | 5.56 d (3.5) | 95.2, $CH_2$ | |
| | 5.36 d (3.5) | | | 5.39 d (3.5) | | |
| 9 | 7.32 d (15.5) | 108.2, CH | 5 | 7.41 d (15.8) | 110.6, CH | 5, 11 |
| 10 | 7.80 d (15.5) | 146.4, CH | | 7.74 d (15.8) | 144.2, CH | 5, 12 |
| 11 | | n.d.[a] | | | 125.3, C | |
| 12 | 6.79 s | 108.0, CH | 10, 13, 14, 16 | 6.70 s | 111.6, CH | 16 |
| 13 | | 143.8, C | | | n.d.[a] | |
| 14 | | 138.2, C | | | n.d.[a] | |
| 15 | | 143.8, C | | 6.70 s | 101.2, CH | 16 |
| 16 | 6.79 s | 108.0, CH | 10, 12, 14, 15 | | 146.2, C | |

[a] Chemical shifts were not detected.

***Table S2.*** Plasmids and strains used in this work

| Bacterial Strain or Plasmid | Relevant Characteristics | Source or Reference |
| --- | --- | --- |
| Bacterial Strains | | |
| *E. coli* | | |
| XL1-blue | cloning strain | Agilent |
| WM6026 | conjugation donor strain, DAP auxotroph | Blodgett et al.[8] |
| BAP-1 | expression strain | Macro Lab UC Berkeley |
| *Csa* | | ATCC 27021 |
| *Csa_empty* | vector background | Herman et al.[6] |
| *Csa_csp* | heterologous expression strain | this study |
| Plasmids | | |
| pJL83 | chloramphenicol/thiamphenicol resistance, *cspD* targeted suicide vector | this study |
| pJL109 | erythromycin resistance, P-bdh *csp* | this study |
| pDG16 | ampicillin resistance | this study |

***Table S3.*** Synthetic oligonucleotides used in this work

| Primer | Sequence (5' to 3') | Target/Role |
|---|---|---|
| JL322 | CATTTGCAGGCTTCTTATTTTTATGGGAAATGTACCTATGGCAGATG | upstream homology region of *cspD* |
| JL323 | CTTGCCCACTGGCCGGCCTCCATTAATTATGCTCAACG | upstream homology region of *cspD* |
| JL324 | GTGTTTTTTGTTACCCTAAGTTTGGATTATCTCAAAGGGAAATAC | downstream homology region of *cspD* |
| JL325 | ATGAGATTATCAAAAAGGAGTTTTTTACTGATTTGATCAACCAC | downstream homology region of *cspD* |
| JL388 | TTACACTATCAAATAATCTATCTATAATCATCTAAGTTCCCTCTCAAATT | *catP* |
| JL389 | TTGAATTTGAGAGGGAACTTAGATGATTATAGATAGATTATTTGATAGTGTAA | *catP* |
| SN7 | GATTGTTATGGATTATAAGCGGCCGGTTCATATTTAT | erythromycin resistance gene |
| SN8 | GGTCATGAGATTATCAAAAAGGAGTTTTAACTTACTTATTAAAT | erythromycin resistance gene |
| JL401 | GTAAGAGGAGGAAAAGAATTGTACATGGAATTGGAATTGAAAAATAAGTTAC | 5' portion of *csp* |
| JL406 | GCCAAAATTCGGTCAAATTTTTACATC | 5' portion of *csp* |
| JL409 | ATGTAAAAATTTGACCGAATTTTGG | center portion of *csp* |
| JL410 | GGAATGTTAAGCATAGCTGAATTATC | center portion of *csp* |
| JL407 | GATAATTCAGCTATGCTTAACATTCC | 3' portion of *csp* |
| JL402 | TAAAACGACGGCCAGTGCCACTAATTATTCTTTCCATTGTTAACAAC | 3' portion of *csp* |
| JL399 | CAGGAAACAGCTATGACCGCGGTAAGACGAACAGCAGAAC | P-bdh promoter of *Csa* |
| JL400 | CTTATTTTTCAATTCCAATTCCATGTACAATTCTTTTCCTCCTCTTACACAC | P-bdh promoter of *Csa* |
| DG28 | GGTTGGGAATTGCAAATGTTTGAAAAGAGAGTAAAAGAAAATCCAGA | 5' portion of A-ACP didomain |
| DG30 | GGAGATGGGAAGTCATTAAATATATTTACTTAAATCTTTTATGCAAAAATTATCA TATAA | 3' portion of A-ACP didomain |

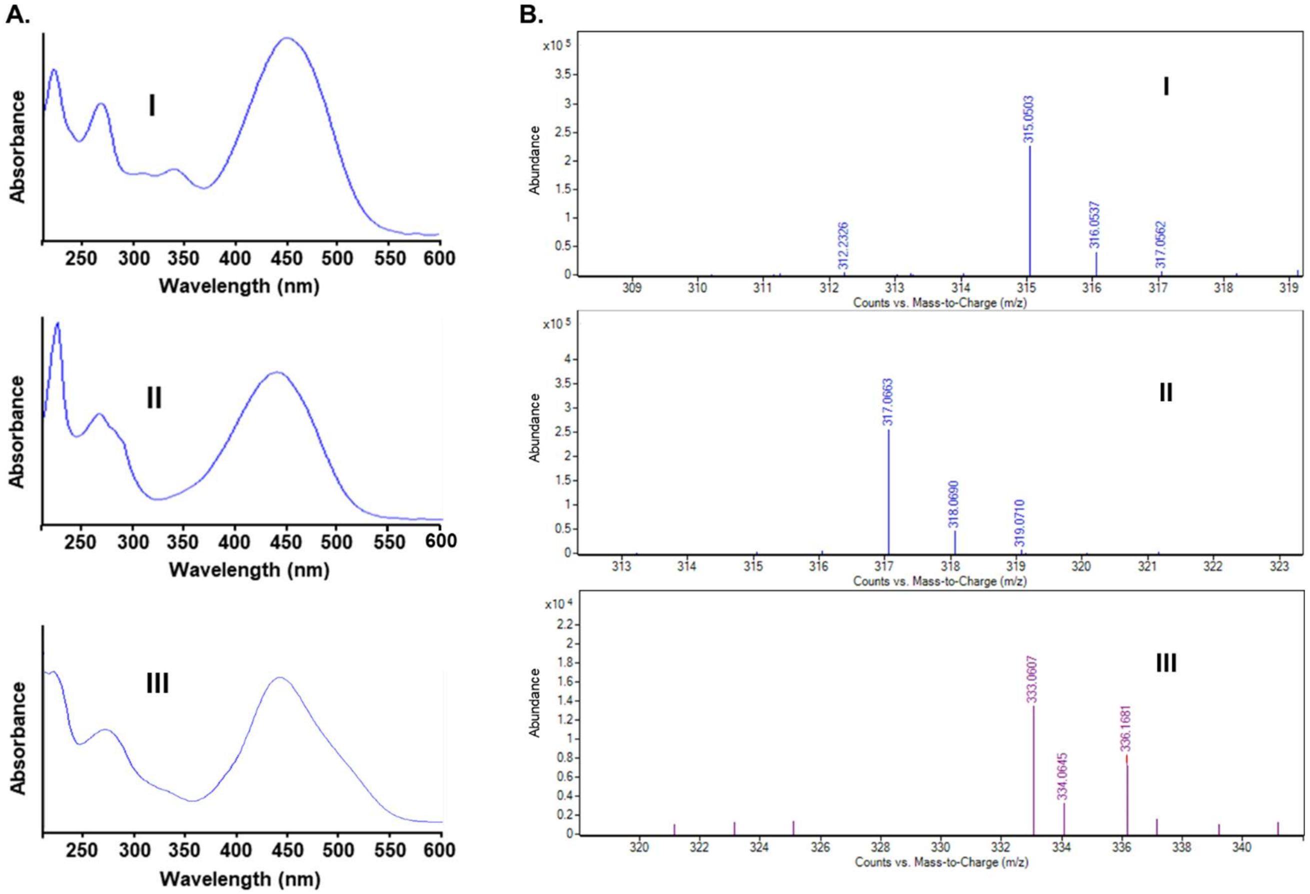


***Figure S1.*** UV and HRMS characterization of clostyrylpyrones. (A) UV spectra of peaks I, II, and III. (B) Positive-mode HRMS spectra of I, II, and III. The different colors are due to the different methods of the mass spec software with different default color settings during obtaining these raw data.

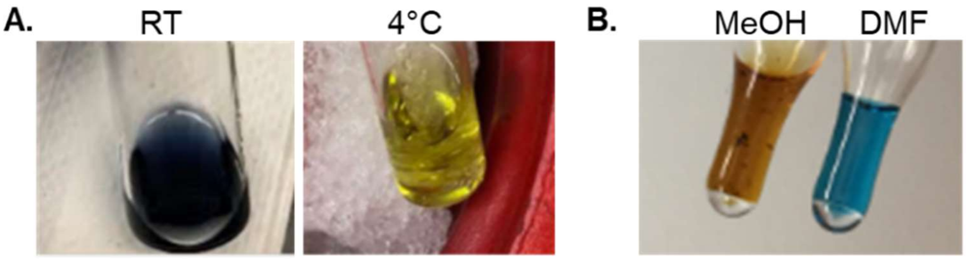


***Figure S2.*** Chromism of **3** and **4**. (A) Thermochromism in 30% acetonitrile in water. (B) Solvatochromism in methanol and dimethylformamide.

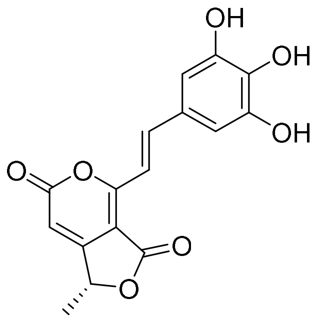


**1**, clostyrylpyrone A

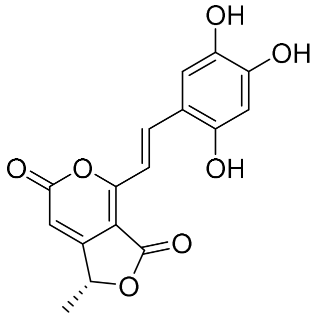


**2**, clostyrylpyrone B

**A.**

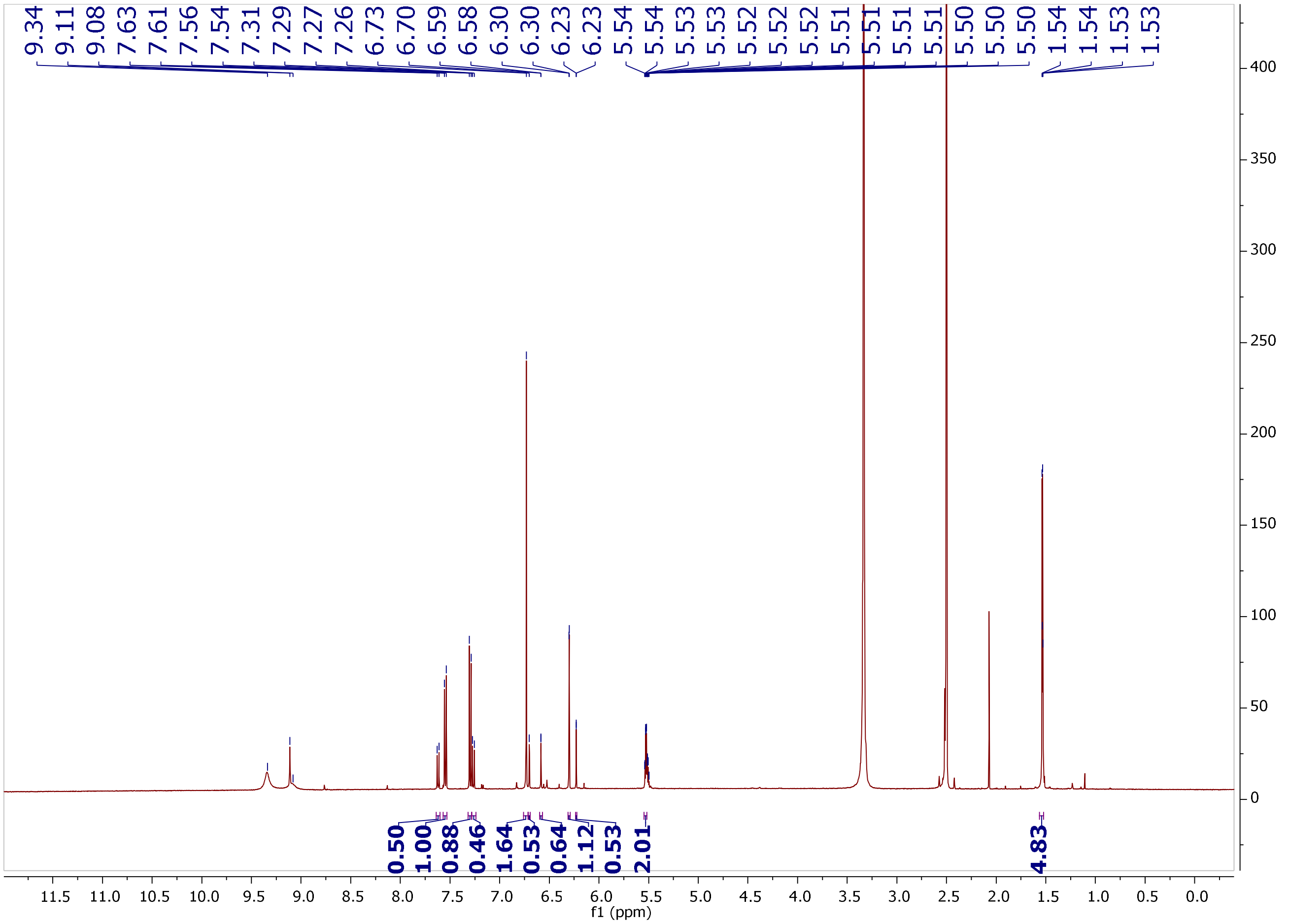


***Figure S3.*** NMR characterization of **1**, **2**. (A) $^{1}$H NMR spectrum of compounds **1**, **2** in DMSO-$d_6$ (900 MHz).

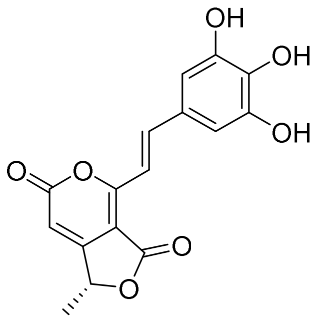


**1**, clostyrylpyrone A

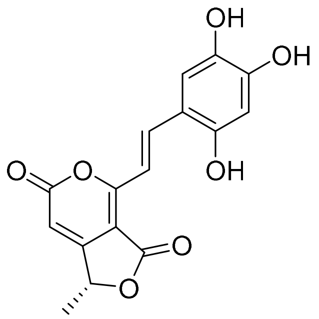


**2**, clostyrylpyrone B

**B.**

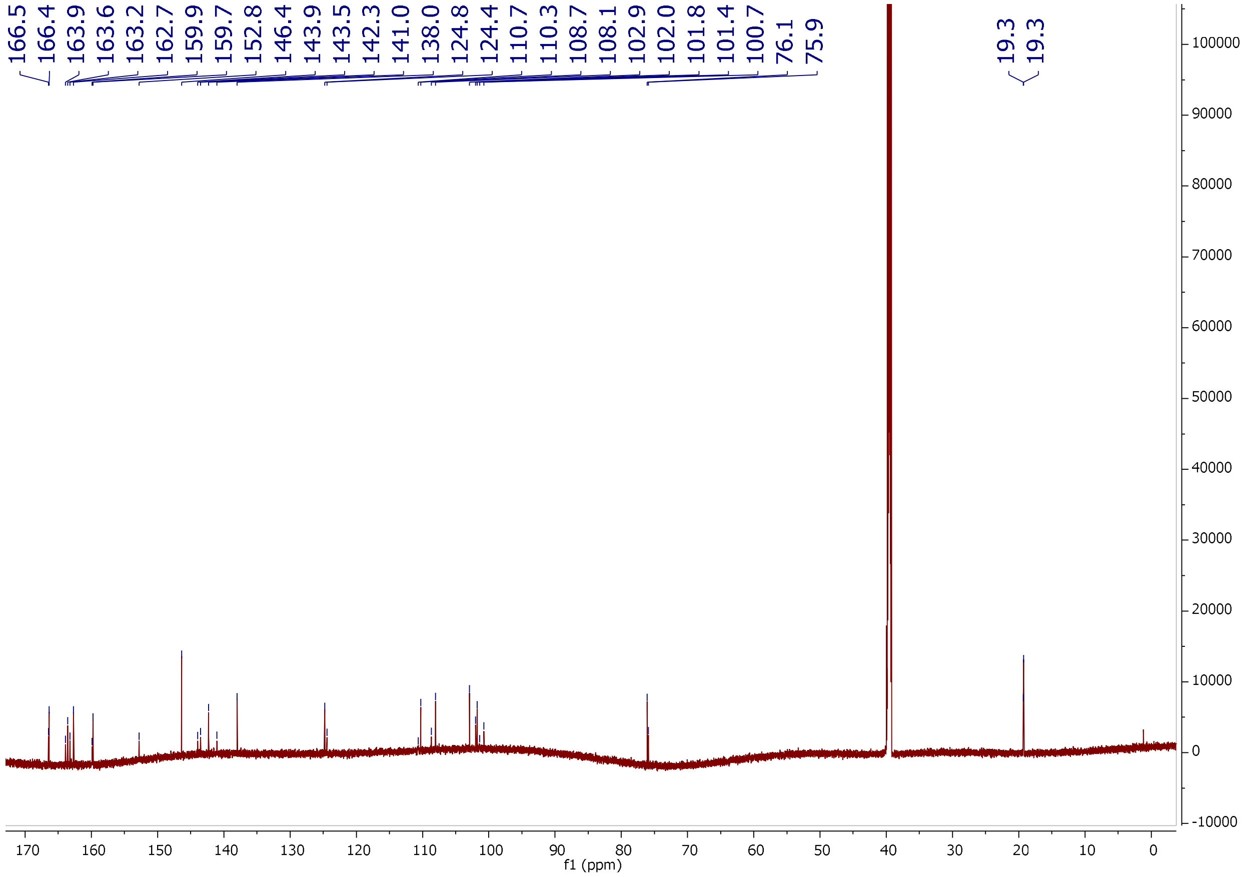


***Figure S3** (continued)*. (B) $^{13}$C NMR spectrum of compounds **1**, **2** in DMSO-$d_6$ (226 MHz).

C.

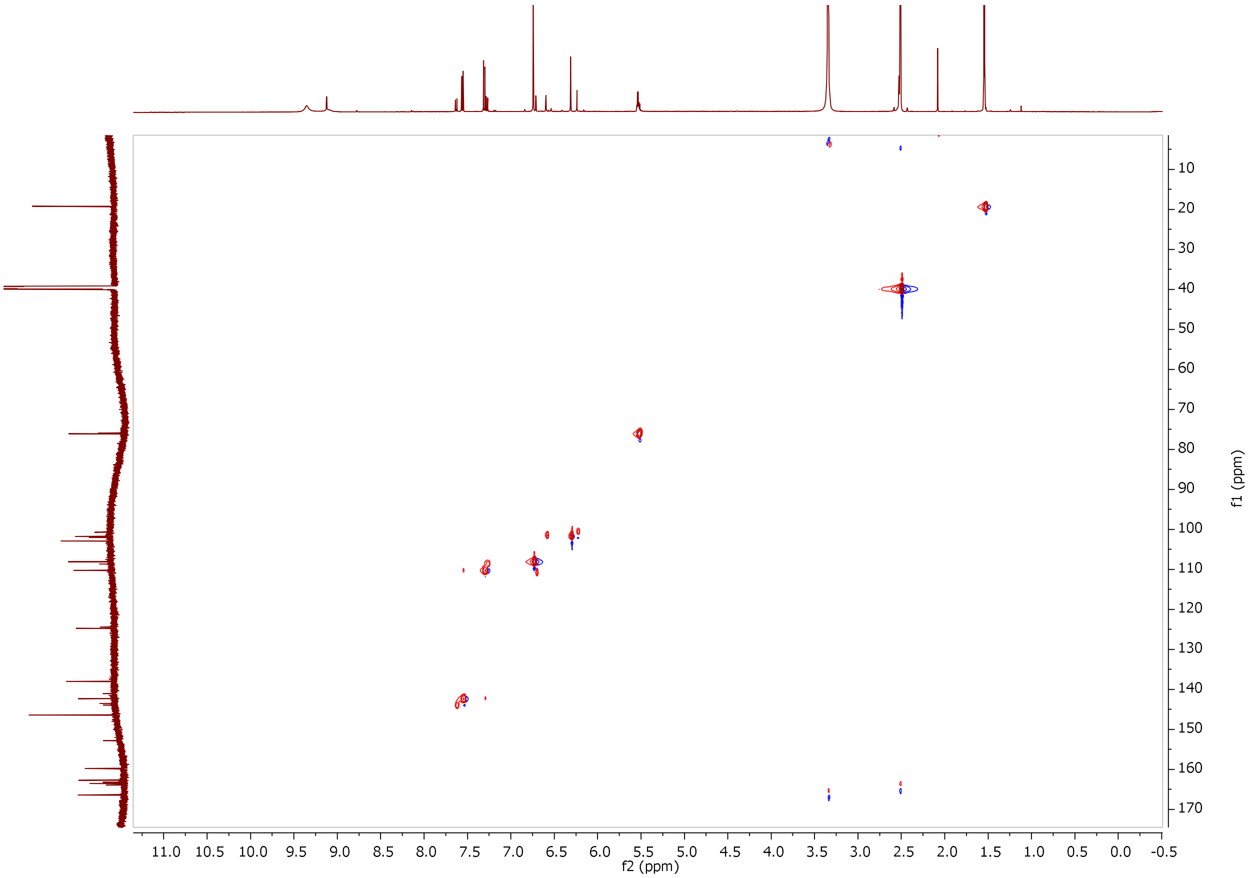


***Figure S3*** *(continued)***.** (C) gHSQC spectrum of compounds **1**, **2** in DMSO-$d_6$ (900 MHz).

D.

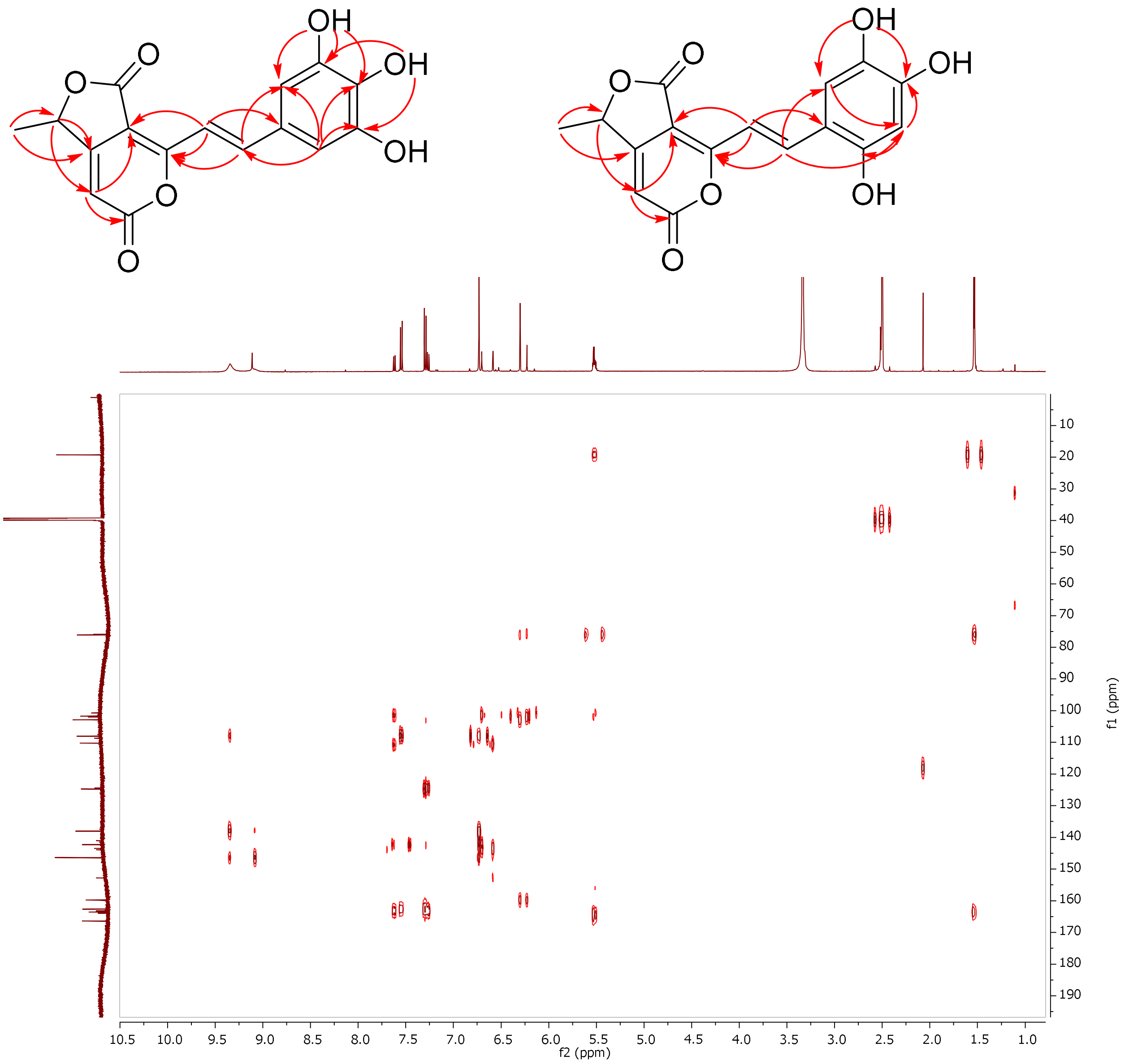


***Figure S3*** *(continued)***.** (D) gHMBC correlations of compounds **1, 2** in DMSO-d$_6$ (900 MHz).

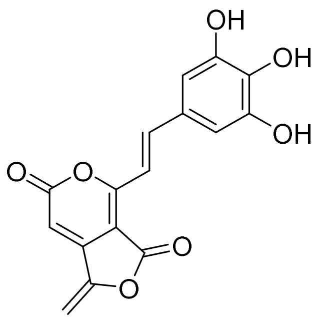


**3**, clostyrylpyrone C

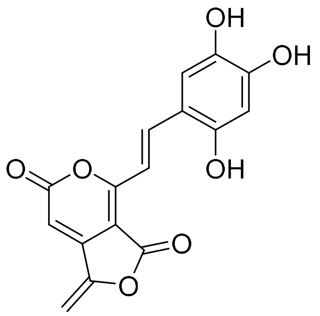


**4**, clostyrylpyrone D

**A.**

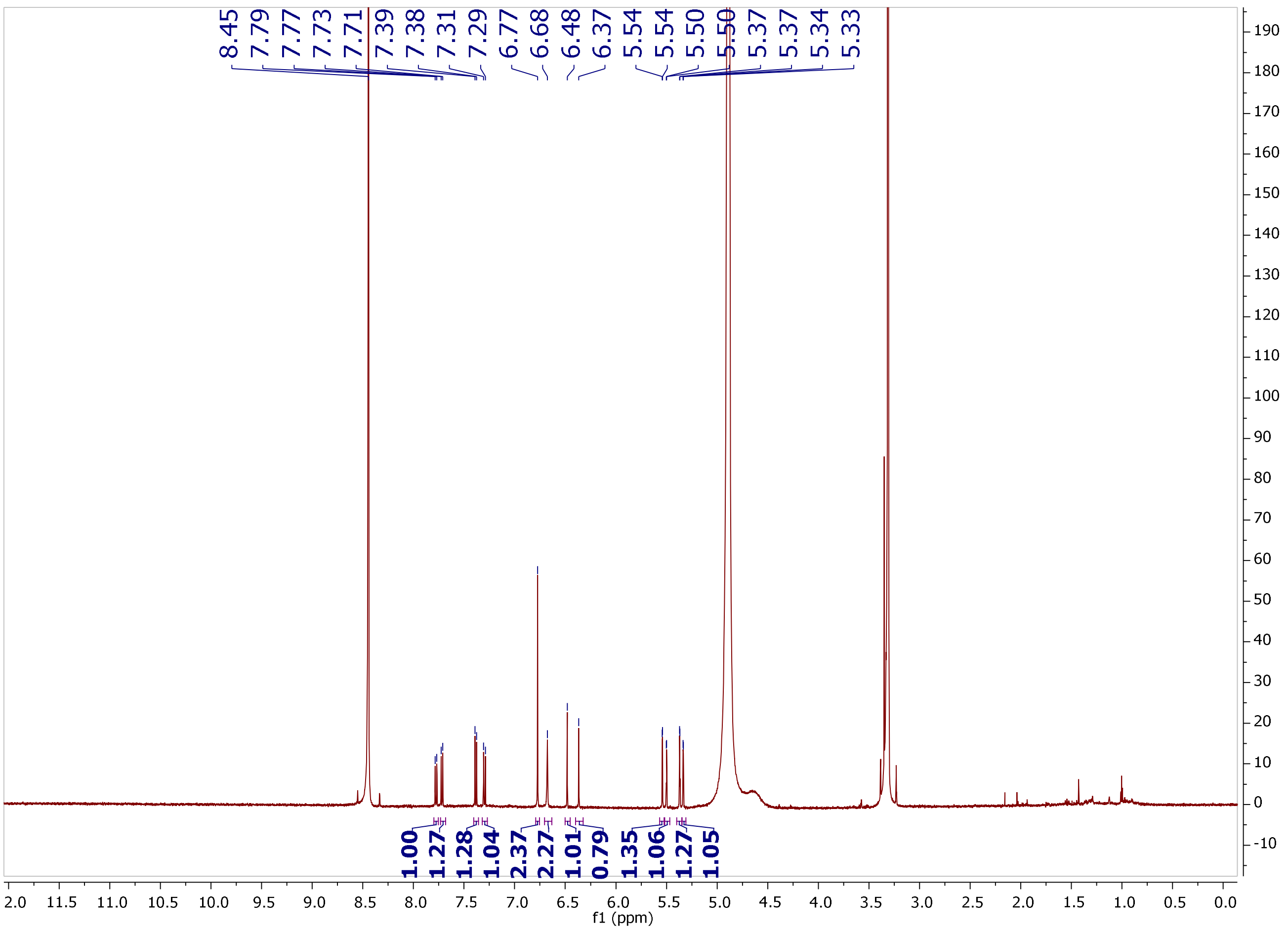


***Figure S4.*** NMR characterization of compounds **3**, **4**. (A) $^{1}$H NMR spectrum of compounds **3**, **4** in $CD_3OD$ (900 MHz).

B.

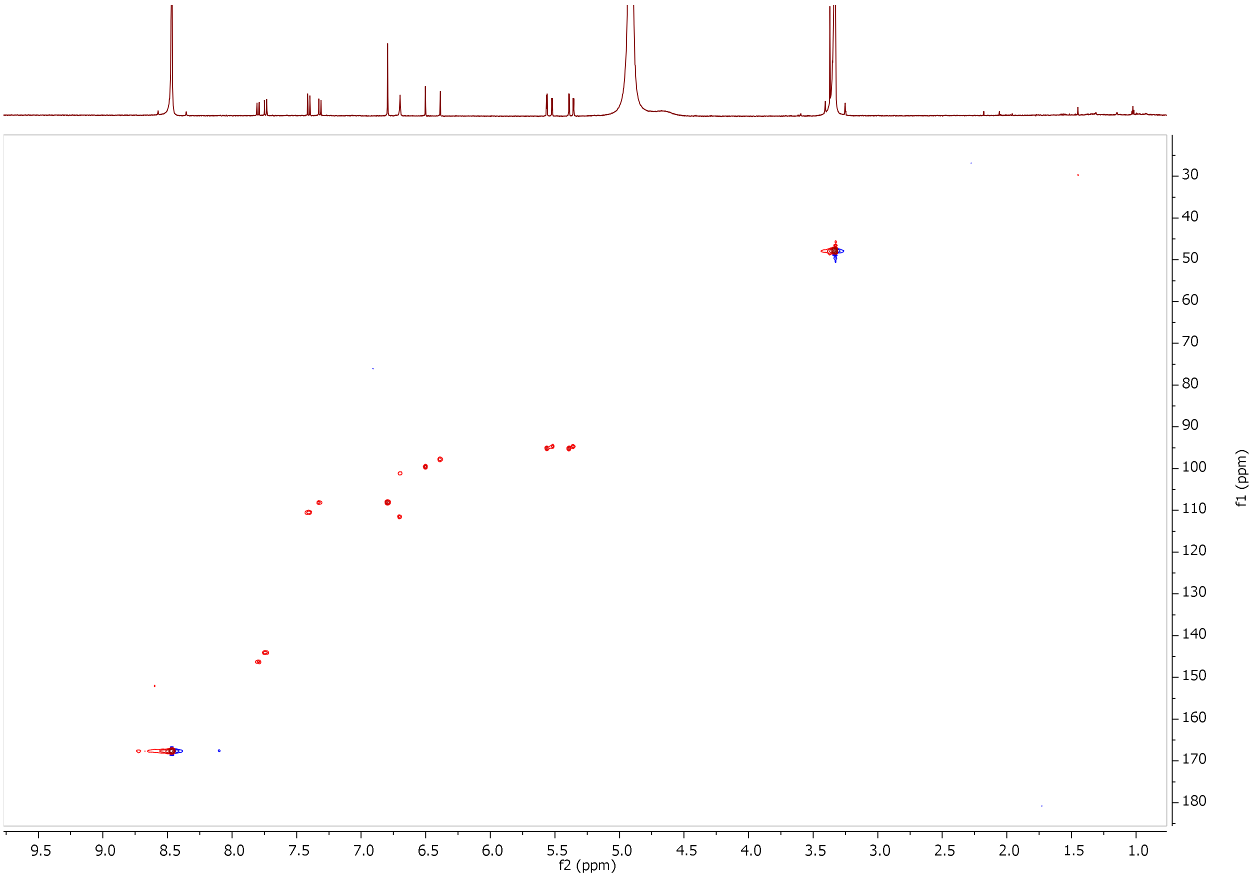


***Figure S4** (continued)***.** (B) gHSQC spectrum of compounds **3**, **4** in $CD_3OD$ (900 MHz).

**c.**

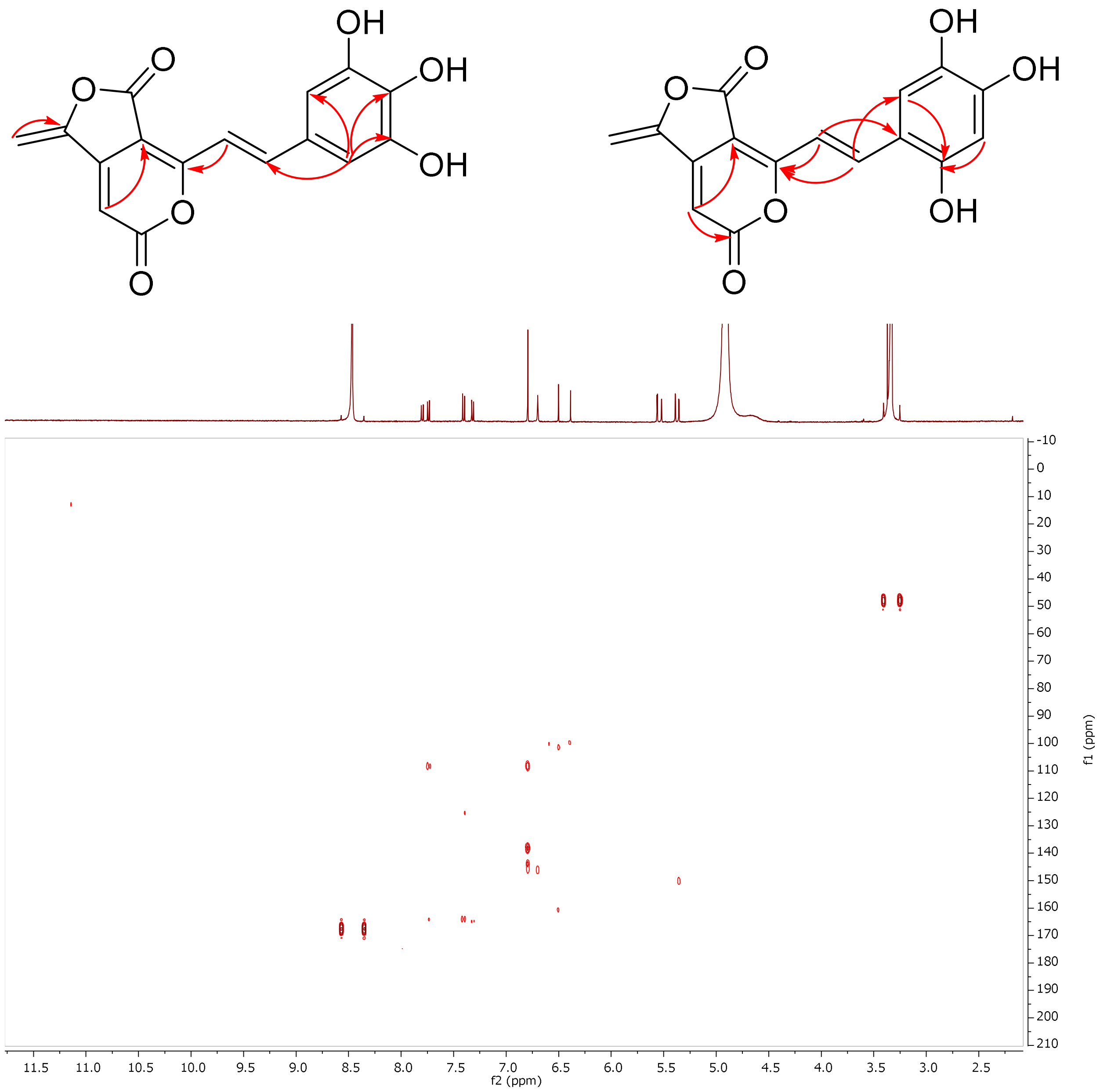


***Figure S4** (continued)***.** (C) gHMBC correlations of compounds **3**, **4** in $CD_3OD$ (900 MHz).

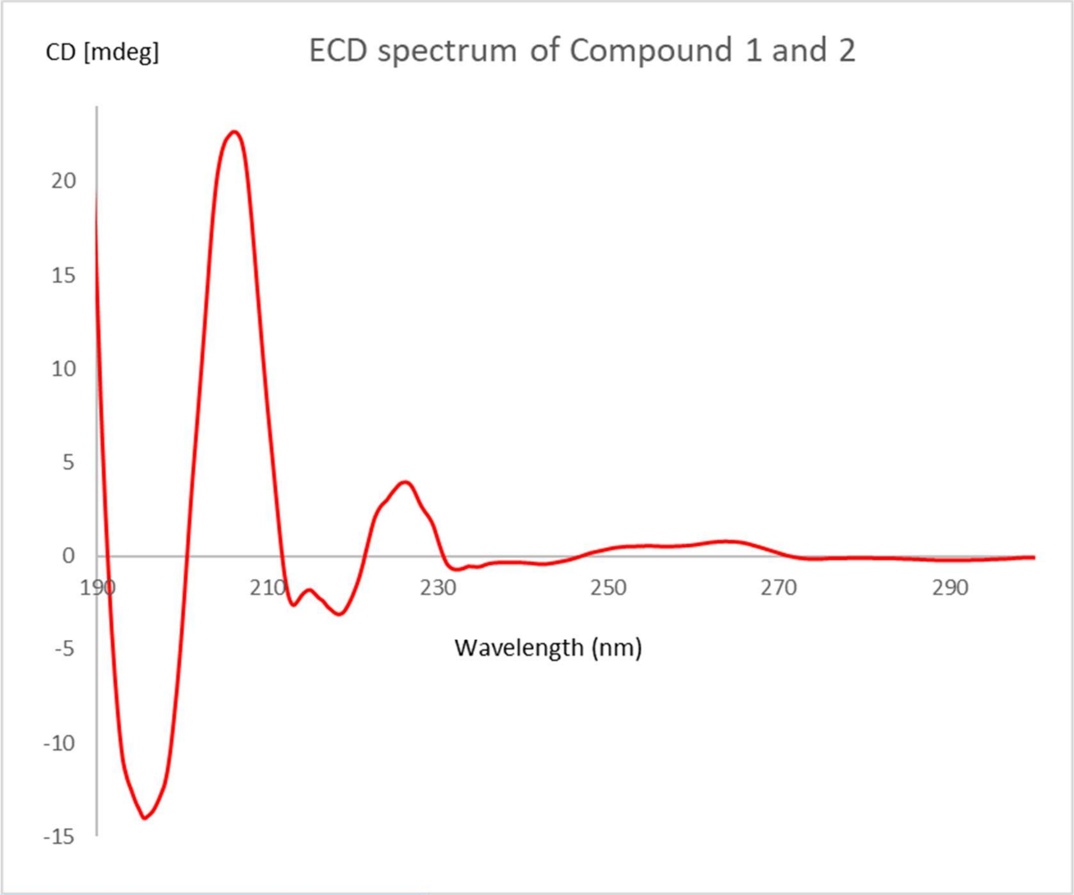


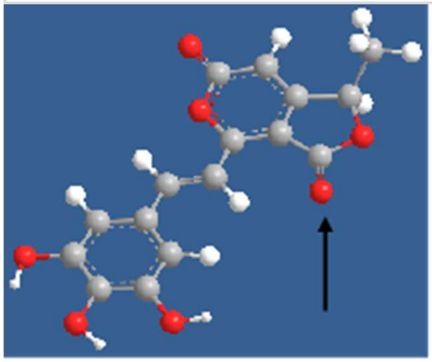

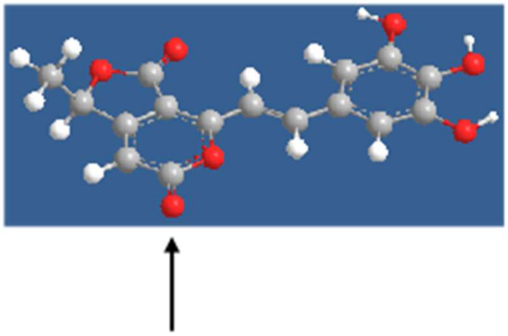

$n \rightarrow \pi^*$
$\Delta\varepsilon_{243\ nm} = -\ 0.40$

$n \rightarrow \pi^*$
$\Delta\varepsilon_{264\ nm} = +\ 0.78$

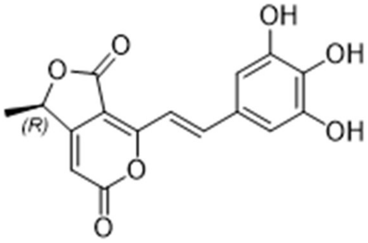


***Figure S5.*** ECD spectrum analysis of compounds **1** and **2.**

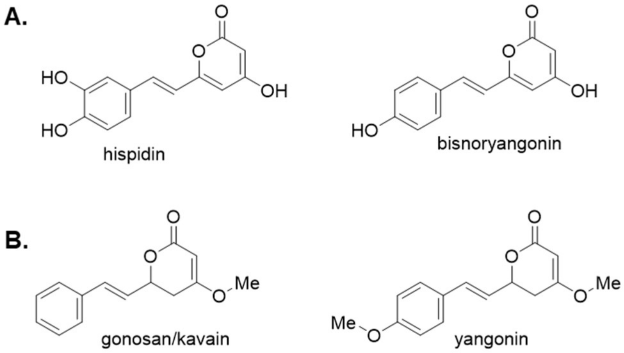


***Figure S6.*** Selected styrylpyrone NPs from (A) fungi and (B) plants.

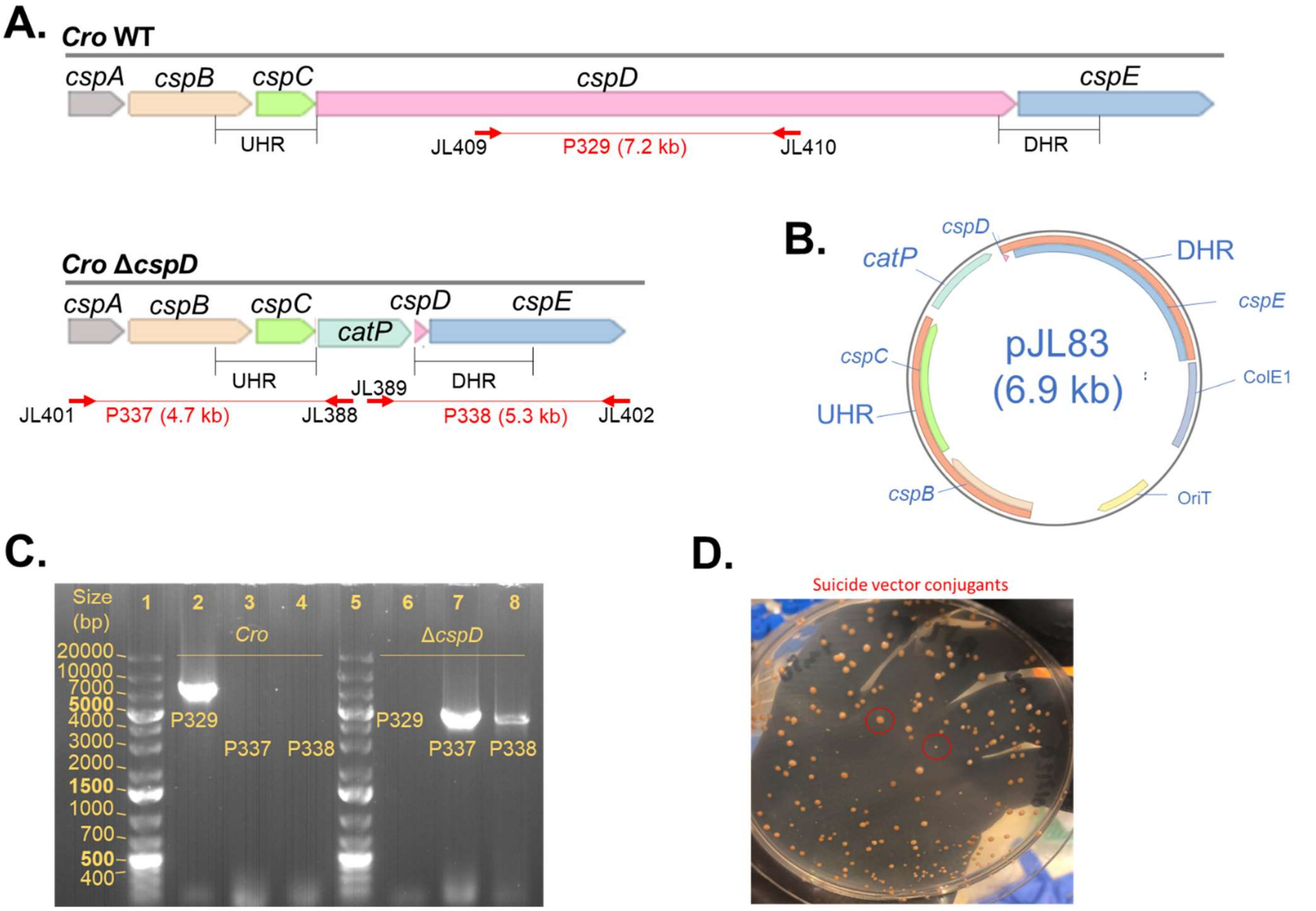


***Figure S7.*** Generation of *Cro* Δ*cspD*. (A) Genotype of *Cro* wild-type and Δ*cspD*. Red arrows and text denote PCR primer binding sites. UHR, upstream homology region; DHR, downstream homology region. (B) Map of suicide vector pJL83. (C) PCR genotyping of isolated mutant. (D) Morphology of conjugant colonies. Representative large and small colonies associated with single- and double crossover mutations, respectively, are indicated by red circles

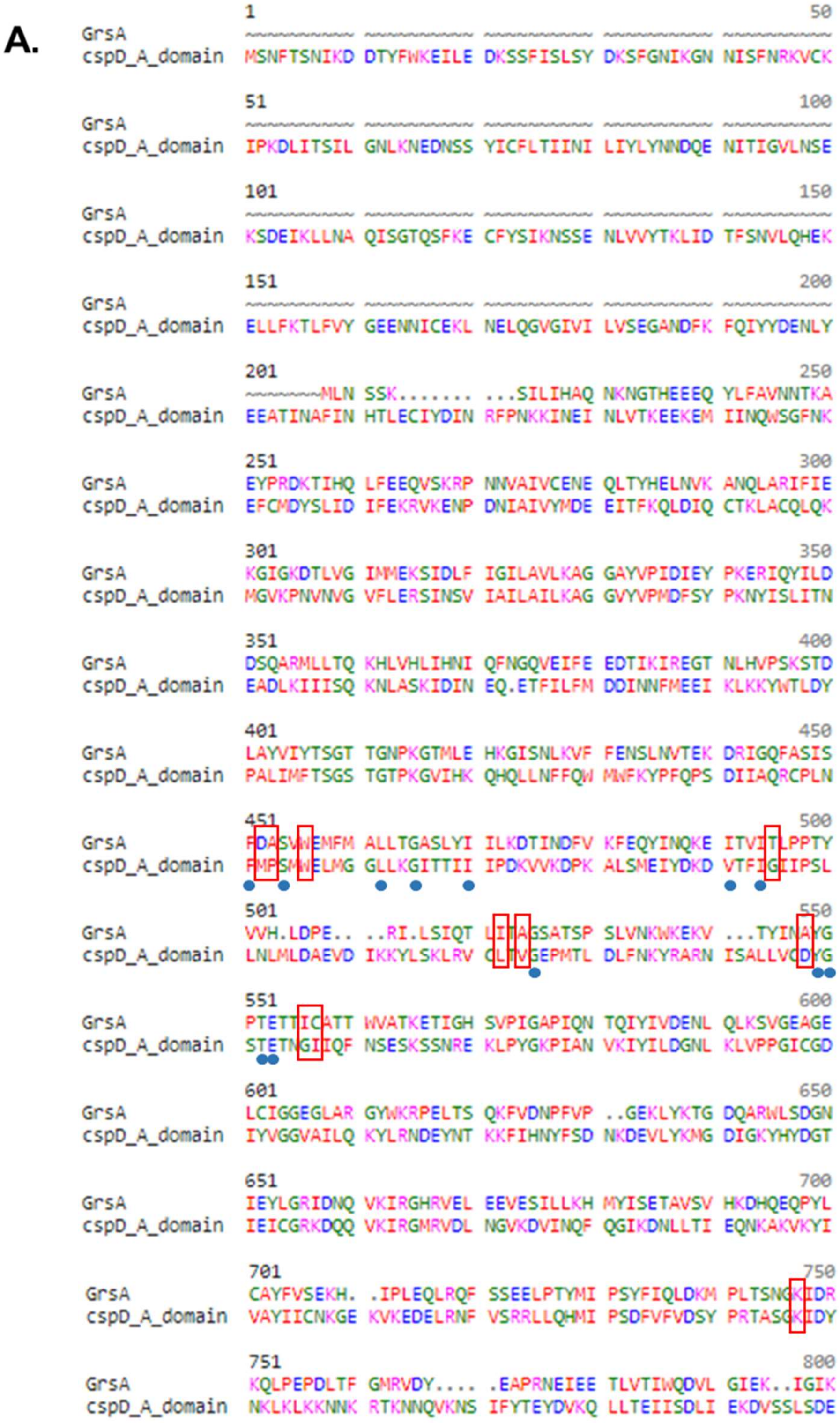


***Figure S8.*** *In silico* analysis of CspD. (A) Protein sequence alignment of GrsA and CspD. Selectivity-conferring residues are boxed in red.[9] Core structural "anchor" residues are indicated with blue dots.[10]

**B.**

| Module | A Domain Signature | | | | | | | | | | Predicted Substrate |
|---|---|---|---|---|---|---|---|---|---|---|---|
| GrsA | 235<br>**D** | 236<br>A | 239<br>**W** | 278<br>**T** | 299<br>I | 301<br>A | 322<br>**A** | 330<br>**I** | 331<br>C | 517<br>**K** | Phe |
| CspD-A | **M** | P | **W** | **G** | L | V | **D** | **G** | I | **K** | No Hit |

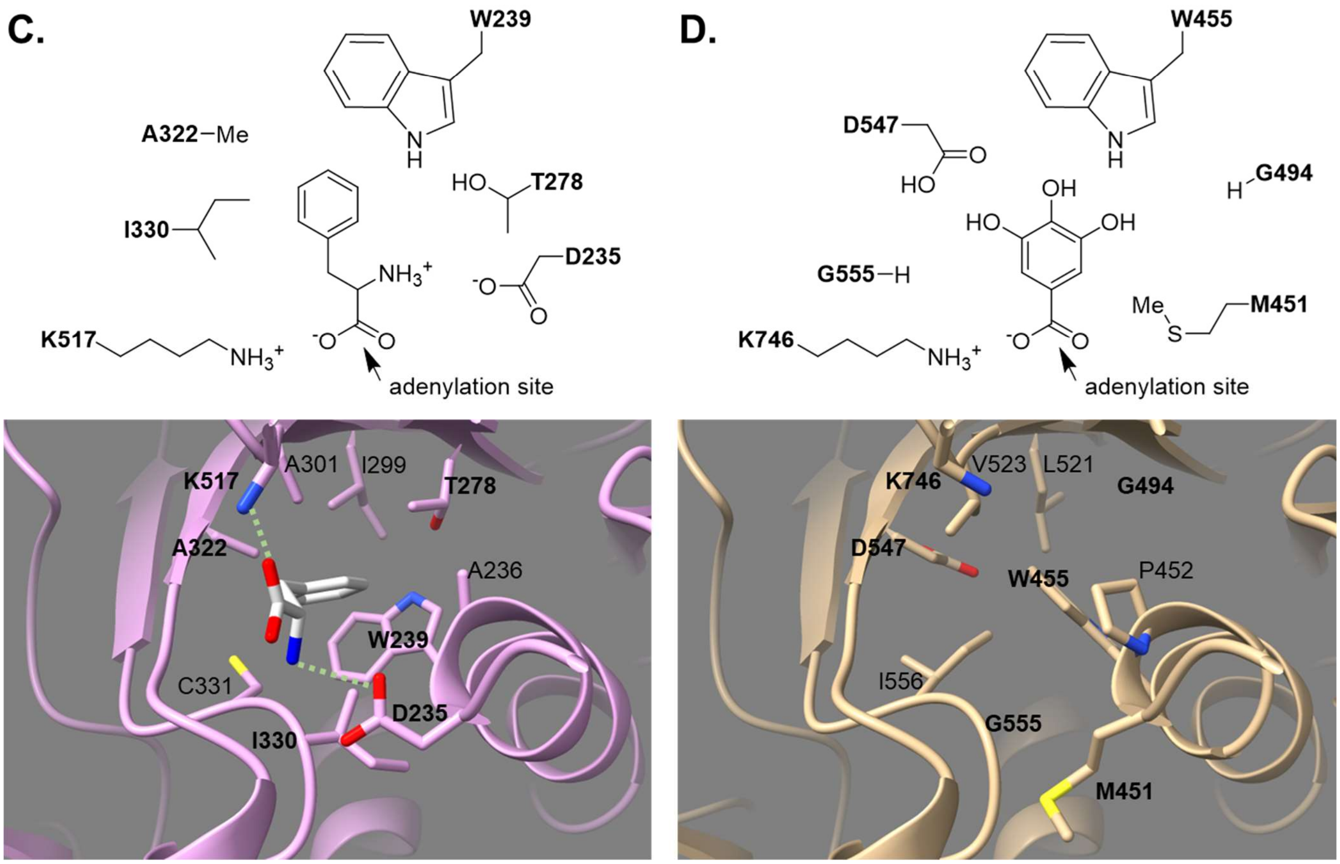


***Figure S8*** *(continued)***.** (B) Selectivity-conferring residues of the A domain of CspD as identified by homology modeling in HHPRED.[11] The corresponding residues in the Phe-activating GrsA (PDB: 1AMU) domain are shown for comparison. (C) Model of the substrate binding pocket of GrsA. Above, relevant residues are illustrated in a simplified depiction. Below, the binding pocket of GrsA is presented a purple ribbon diagram. The Phe substrate is depicted as white sticks. Heteroatoms in Phe or the adjacent side chains are colored as follows: blue, nitrogen; red, oxygen; yellow, sulfur. Green dashed lines indicate relevant interactions. (D) Model of the A domain substrate binding pocket of CspD. Above, analogous specificity residues are illustrated, with gallic acid as a representative substrate. CspD retains the conserved Lys for bonding the carboxylic acid but lacks the Asp residue to interact with a substrate amino group. The substrate specificity may also be affected by the difference in hydrogen-bonding environment of the aromatic pocket. The structural model was obtained by threading CspD, truncated to 959 residues, onto GrsA using MODELLER.[12] Results were visualized in ChimeraX.[13]

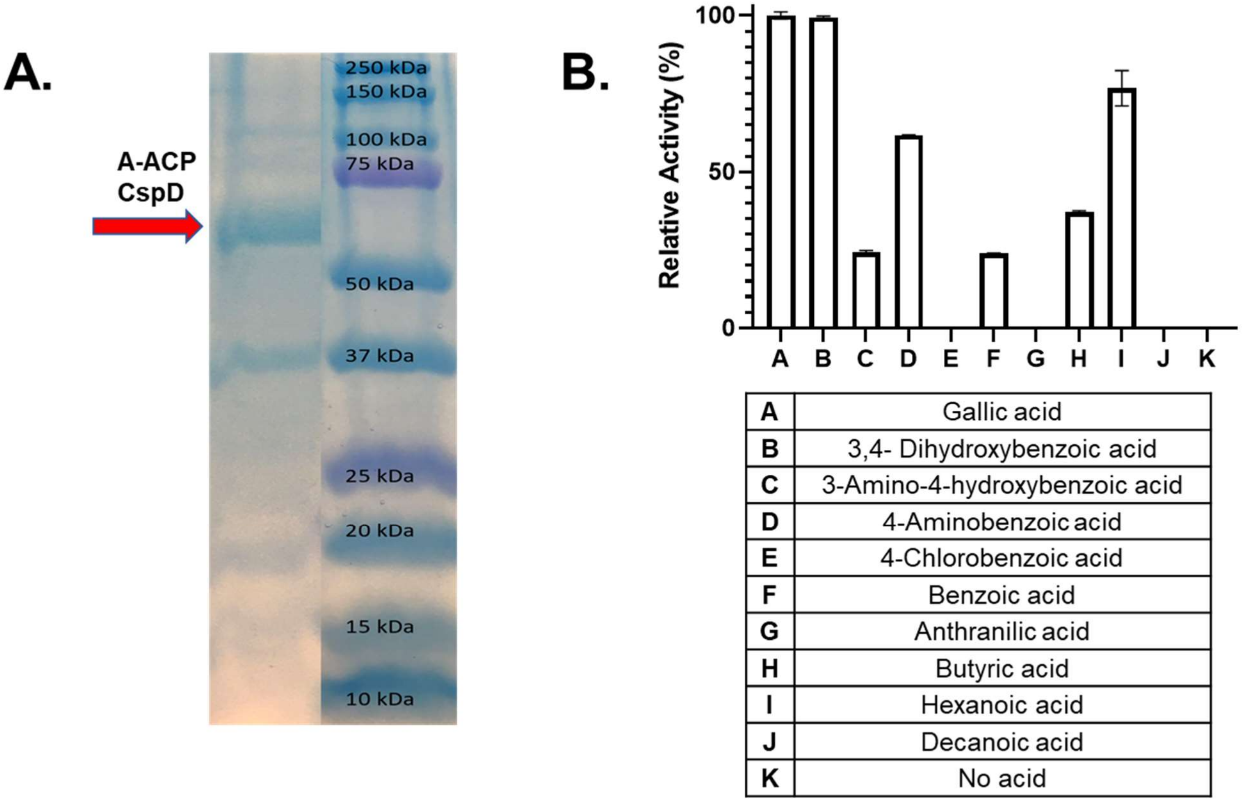


| | |
|---|---|
| A | Gallic acid |
| B | 3,4- Dihydroxybenzoic acid |
| C | 3-Amino-4-hydroxybenzoic acid |
| D | 4-Aminobenzoic acid |
| E | 4-Chlorobenzoic acid |
| F | Benzoic acid |
| G | Anthranilic acid |
| H | Butyric acid |
| I | Hexanoic acid |
| J | Decanoic acid |
| K | No acid |

***Figure S9.*** Biochemical analysis of A-ACP didomain of CspD. (A) SDS-PAGE analysis of recombinant A-ACP didomain of CspD. The A-ACP didomain of CspD with an N-hexahistidine tag was purified from *E. coli* and used for *in vitro* biochemical assays with different acid substrates. Any kD Mini-PROTEAN TGX gels (precast, Biorad) were used for analysis and demonstrated the protein is largely soluble in *E. coli* (67.1 kDa). (B) Relative activity of the A-ACP didomain of CspD towards gallic acid and other substrates (normalized with respect to gallic acid activity).

A.

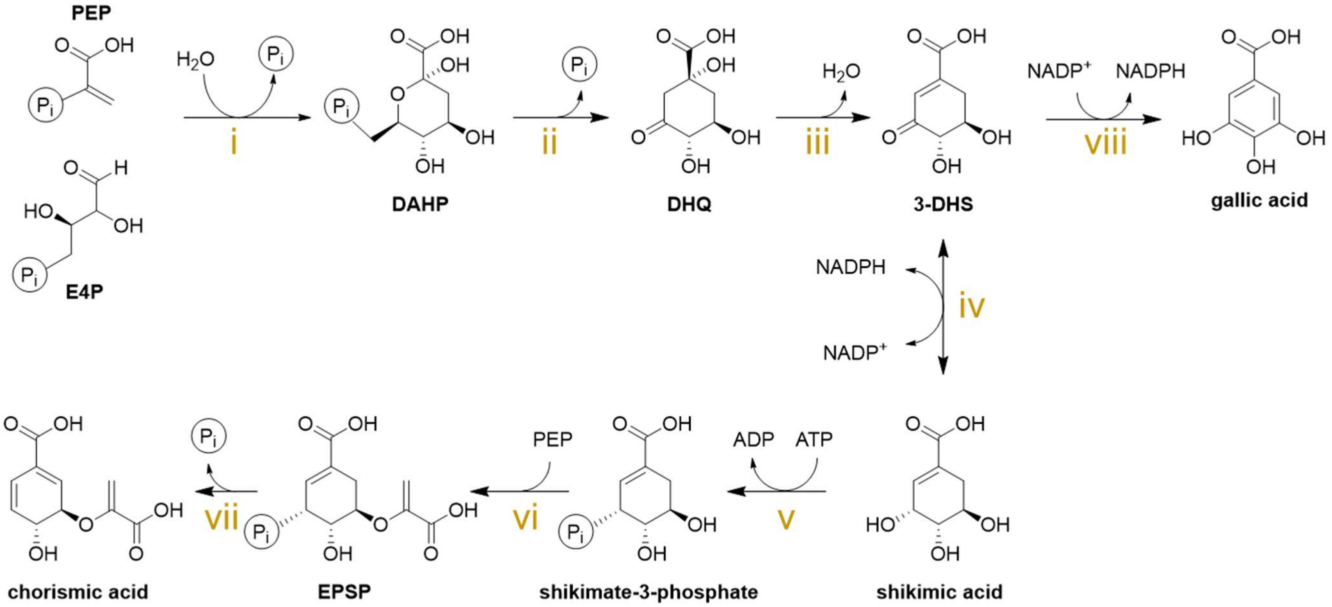


| Step | Putative Function | *Cac* | | Identity/ Similarity (%/%) | *Cro* | |
|---|---|---|---|---|---|---|
| | | Homolog | Locus Tag | | Homolog | Accession |
| i | DAHP synthase | *aroF* | WP_010964210.1 | 96/98 | *aroF* | OOM02945.1 |
| | | | | 25/44 | *aroF* | OOM01708.1 |
| ii | DHQ synthase | *aroB* | WP_010964212.1 | 79/91 | *aroB* | OOM02943.1 |
| iii | type I DHQ dehydratase | | | - | *aroD* | OOM03356.1 |
| | type II DHQ dehydratase | *aroQ* | WP_010964217.1 | - | | |
| iv | dehydrogenase | *aroE* | WP_010964215.1 | 70/86 | *aroE* | OOM02940.1 |
| viii | | | | 32/51 | *aroE* | OOM03357.1 |
| V | shikimate kinase | *aroK* | WP_010964216.1 | 69/85 | *aroK* | OOM02939.1 |
| Vi | EPSP synthase | *aroA* | WP_010964213.1 | 79/90 | *aroA* | OOM02942.1 |
| Vii | chorismate synthase | *aroC* | WP_010964214.1 | 88/93 | *aroC* | OOM02941.1 |

B.

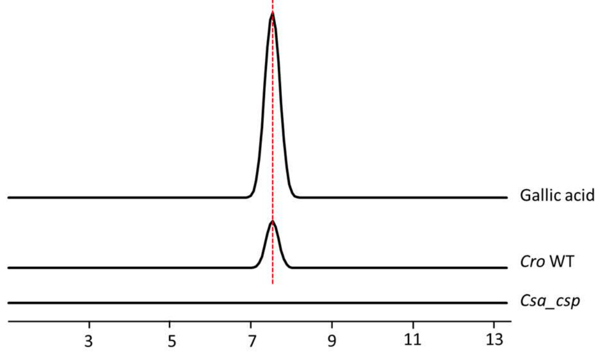


***Figure S10.*** Proposed pathway of gallic acid biosynthesis in *Cro.* (A) The pathway derives from aryl acid biosynthesis, which is found in closely related organisms such as *Clostridium acetobutylicum* (*Cac*). Abbreviations: DAHP, 3-deoxy-D-arabinoheptulosonate 7-phosphate; DHQ, 3-dehydroquinate; 3-DHS, 3-dehydroshikimate; EPSP, 5-enolpyruvylshikimate-3-phosphate. (B) LC-HRMS analysis of metabolites. Extracted ion chromatograms are shown. The calculated mass of gallic acid (m/z = 171.0288 $[M+H]^+$) with a 10-p.p.m. mass error tolerance was used. *Cro* WT can produce gallic acid while *Csa_csp* cannot.

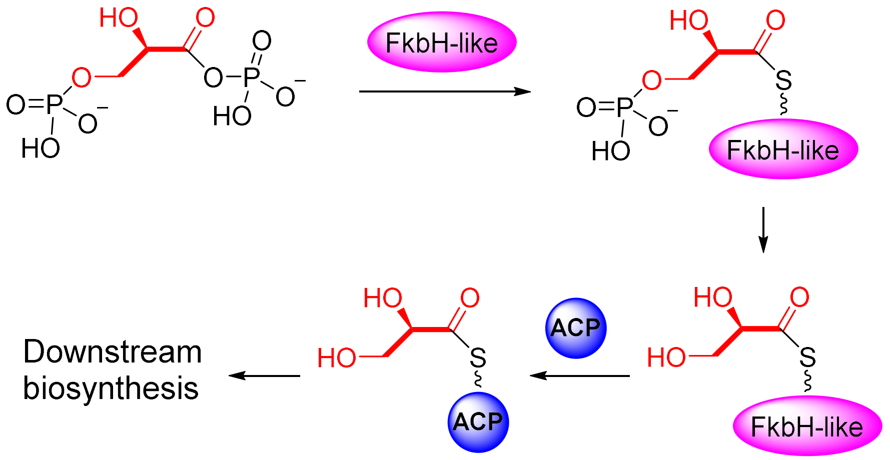


***Figure S11.*** The activity of FkbH-like domain. A typical FkbH-like domain can divert the primary metabolite 1,3-diphosphoglycerate into the polyketide biosynthetic pathway via glycerate.

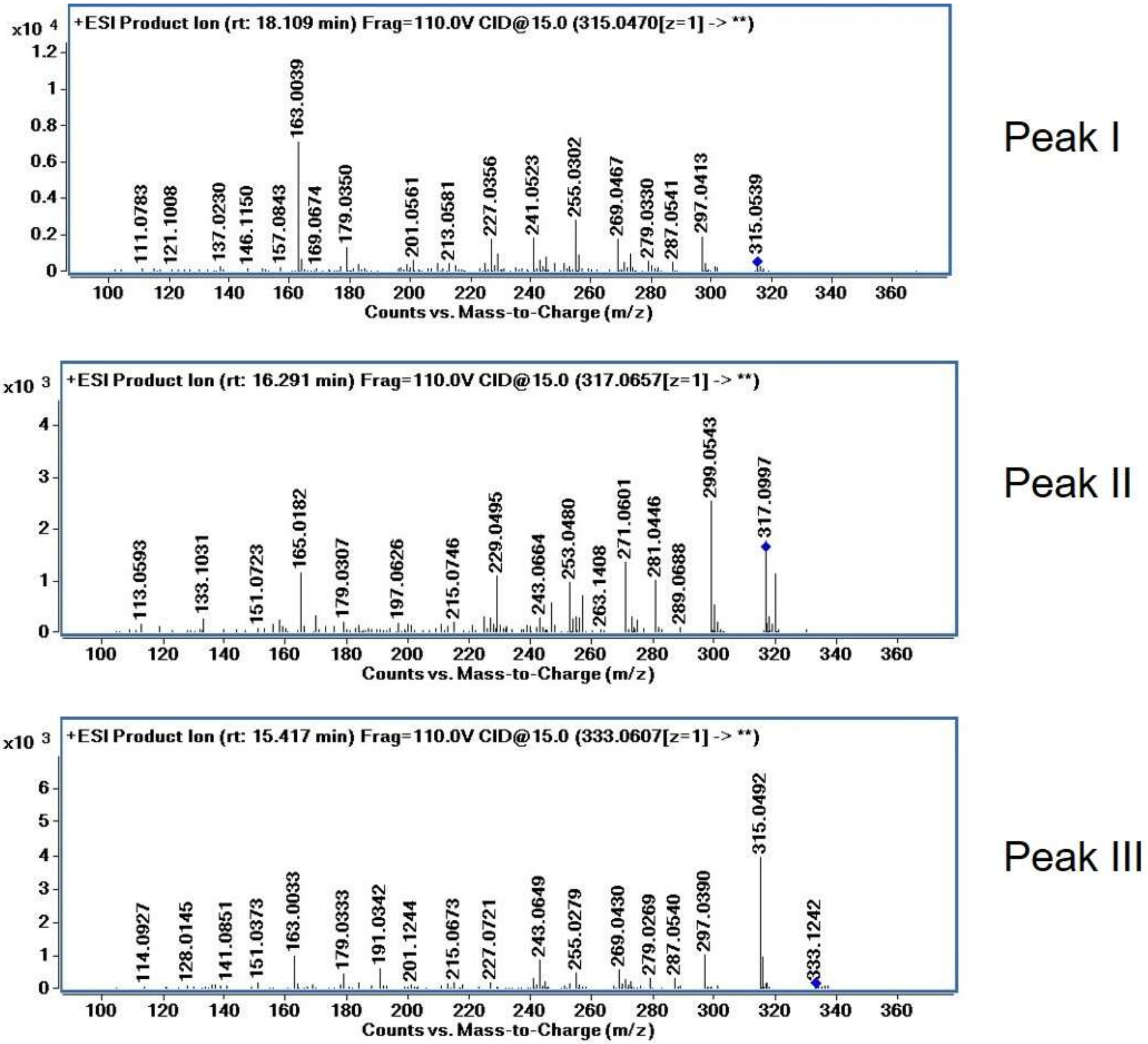


***Figure S12.*** MS/MS spectra of peak I, II, III.

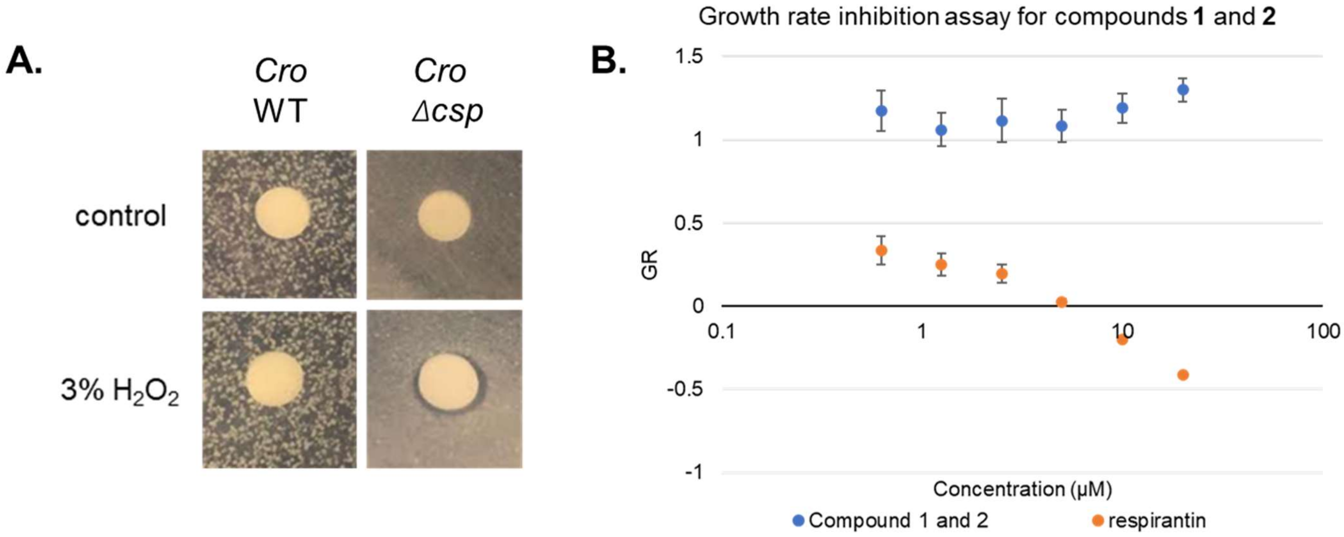


***Figure S13.*** Bioactivity assays of clostyrylpyrones. (A) Hydrogen peroxide disc diffusion assays. A slight inhibition zone is visible in the *Cro* mutant deficient in clostyrylpyrone production. (B) Growth rate (GR) inhibition assay of compound **1** and **2**. The GR value ranges from -1 to 0 for cell death, and 0 to 1 for growth rate inhibition. Inhibition is detected in the positive control, performed with respirantin, but not in compounds **1** and **2**.
